# Secretome characterization of the lignocellulose-degrading fungi *Pycnoporus sanguineus* and *Ganoderma applanatum*

**DOI:** 10.1101/2020.09.22.308908

**Authors:** Albertina Gauna, Alvaro S. Larran, Susana R. Feldman, Hugo R. Permingeat, Valeria E. Perotti

## Abstract

C4 grasses are common species in rangelands around the world and represent an attractive option for second-generation biofuels production. Although they display a high polysaccharide content and reach great levels of biomass accumulation, there is a major technical issue to be solved before they can be considered as biofuels feedstock: lignin removal. Concerning this, *Pycnoporus* and *Ganoderma* fungal genera have been highlighted due to their ability to hydrolyze lignocellulose. The goals here were to evaluate the pretreatment efficiency using *P. sanguineus* and *G. applanatum* secretomes harvested from a glucose-free inductive medium and to identify the fungal enzymatic activities responsible for the lignin degradation and glucose release. The findings show that *P. sanguineus* secretome exhibits a higher activity of lignocellulolytic enzymes compared to the one from *G. applanatum*. Interestingly, zymograms in presence of glucose suggest that a β-glucosidase isoform from *P. sanguineus* could be glucose-tolerant. The proteomic approach carried out allowed to identify 73 and 180 different proteins for *G. applanatum* and *P. sanguineus* secretomes, respectively, which were functionally classified in five main categories, and a miscellaneous group. Many uncharacterized proteins were found in both secretomes, reflecting that greater research is still needed for a better comprehension of lignocellulose degradation.

## 1. Introduction

Energy demand has grown exponentially since the Industrial Revolution, with the collateral effect of increasing CO_2_ and other greenhouse gases levels in the atmosphere. Intergovernmental Panel on Climate Change (IPCC 2014) claims these gases are responsible for global climate change. Therefore, during the last decade, there has been a rising interest in renewable energy sources, especially in second generation biofuels, which do not compete with food supplies. In this context, different rangeland C4 grasses have been proposed as potential sources of fermentable sugars (Limayem and Ricke, 2012; Alfenore and Molina-Jouve, 2016).

*Panicum prionitis* Ness (=*Coleataenia prionitis* (Ness) Soreng), a dominant C4 grass species in plant communities of the floodplain and islands of the Paraná River (Argentina), represents an attractive option for bioethanol production. *P. prionitis* can reach high levels of biomass accumulation (22.35 Mg of dry matter per hectare; Sosa et al. 2019) and displays a high polysaccharide composition (around 70% of dry weight, as reported in Gauna et al., 2018). However, there is a major technical issue to be addressed before it can be used for bioethanol industrial production: lignin removal. Briefly, lignocellulosic biomass can be converted into ethanol in four steps: pretreatment, enzymatic hydrolysis, fermentation, and distillation of ethanol produced (Salehi Jouzani and Taherzadeh, 2015; Robak and Balcerek, 2018). During pretreatment, lignin within cell wall (about 7% in *P. prionitis*) must be removed to allow the access of hydrolytic enzymes to cellulose and hemicellulose. This procedure is usually the bottleneck for economically viable second-generation biofuel production (Mood et al., 2013; Masran et al., 2016; Cheah et al., 2020).

Few microorganisms are capable of hydrolyzing lignocellulose. Among them, basidiomycetes causing white rot wood decay are particularly effective because they use lignocellulose of plant cell walls as a carbon source during secondary metabolism triggered by nutrient starvation (Chandel et al., 2015). These microorganisms synthesize and secrete a considerable number of hydrolytic enzymes, including cellulases, hemicellulases, lignin-modifying enzymes, and other accessory enzymes (Manavalan et al., 2015) which can be employed in a wide range of industrial processes (Camarero et al., 2002; Georis et al., 2003; Polizeli et al., 2005).

Some species of *Pycnoporus* and *Ganoderma* genera have been highlighted due to their ability to synthesize valuable molecules including antioxidants, antibiotics, anti-inflammatory, and anti-tumoral compounds, as well as to efficiently produce laccases, xylanases, and other industrial interesting enzymes (Smânia et al., 2003; Zhou et al., 2012; Yu et al., 2015). White rot basidiomycetes from *Pycnoporus* spp. are widely studied for their ability to: (i) secrete thermostable laccases (Eugenio et al., 2009; Lu et al., 2010; Uzan et al., 2010) and endoxylanases (Niderhaus et al., 2018) as part of their ligninolytic system, (ii) produce efficient cellulolytic and hemicellulolytic extracts (Scarpa et al., 2019), and (iii) produce α-amilases (Siqueira et al., 1997).

Current advances in MS-based proteomics provide powerful tools for studies that involve complex biological systems, even more after the increasing number of genomes from filamentous fungi that have become recently available (Guo et al., 2018; Shankar et al., 2019; Jain et al., 2020).

In previous works, the group has assessed the potential of *P. sanguineus* and *G. applanatum* secretomes (extracellular proteome) to act as efficient biological pretreatment agents in lignin degradation and glucose release, using green and senescent biomass of *Spartina argentinensis* Parodi (*=Sporobolus spartinus* (Trin.) P.M. Peterson and Saarela) (Larran et al., 2015) and *P. prionitis* (Gauna et al., 2018). The goals here were to evaluate the pretreatment efficiency using secretomes harvested from an inductive medium (using *P. prionitis* leaves as a carbon source) and to identify the fungal enzymatic activities responsible for such lignin degradation and glucose release. A proteomic approach was carried out to promote a deep characterization of the secretomes from these filamentous fungi.

## 2. Materials and methods

### 2.1 Biomass, chemicals and enzymes

*P. prionitis* plants were originally collected in the flooding valley of the Parana River in Argentina (32°52′43.04″S; 60°35′0.33″W) and were treated as described in Gauna et al. (2018). Chemicals and enzymes used were purchased from Sigma-Aldrich Argentina unless stated otherwise.

### 2.2 Fungal species

Two white rot fungi were assessed: *P. sanguineus* and *G. applanatum*. Fungal basidiocarps were collected as described in Gauna et al. (2018), and stock cultures were maintained on Potato Glucose Agar (PGA) at 4°C in the dark.

For inoculation with fungal spores, a 3 mm^3^ plug was transferred to the center of a PGA-containing Petri plate and grown at 28 °C for 7 days. Fungal spores were obtained by washing Petri dishes with 10 mL of 8 g/L sodium chloride supplemented with 40 μL of Tween 20, and then filtering the supernatant through a sterile paper filter to remove the mycelia. Spores were quantified using a Neubaüer counting chamber and were maintained in 20% glycerol at −80 °C.

### 2.3 Pretreatments with fungal secretomes

A 3 mm^3^ piece from the peripheral region of each fungus actively growing on PGA plates was inoculated into flasks containing 20 mL of 50 mM sterile sodium acetate buffer pH 6 with *P. prionitis* ground leaves (50-50% senescent-green) at 7.5 % (w/v) (Inductive Medium, IM). Flasks were incubated at 28 °C with rotary shaking (120 rpm) for 7 days. Secretomes were filtered under sterile conditions and used as pretreatment agents.

Erlenmeyer flasks containing 50 mg of ground dried green or senescent leaves of *P. prionitis* were incubated at 37 °C for 48 h, after the addition of 0.5 mL of fungal secretome diluted in 4.5 mL of 50 mM sodium acetate buffer pH 4. Pretreatments were performed over three biological replicates of *P. prionitis* biomass and were followed by a hydrolysis step. Blanks and controls were prepared as detailed in Gauna et al. (2018). After the whole process, a 20 µL aliquot was taken for glucose determination using an enzymatic glycemia kit (Wiener Lab, Rosario, Argentina).

Hydrolyzed cellulose and the contribution of each step to the total glucose released were calculated as described previously (Larran et al., 2015).

### 2.4 Plate assays and zymogram analysis

*P. sanguineus* and *G. applanatum* were grown in two conditions: IM described in the section 2.3 and Wheat Bran (WB) medium, using this alternative carbon source at 7.5% (w/v) in 50 mM sodium acetate buffer pH 6. A suspension of 1.10^6^ spores was used to inoculate 100 mL of both media in 500 mL baffled flasks. After 7 days, secretomes were used to assay cellulase, xylanase, manganese peroxidase (MnP), and laccase activities in Petri dishes. Negative controls were prepared by boiling the fungal secretomes for 5 minutes.

Protein concentration from each secretome was estimated using the Bio-Rad protein assay reagent (Bio-Rad, Hercules, CA, USA) and bovine serum albumin as standard. Twenty μg of total secreted proteins were added into different wells within each plate and incubated overnight at 30 °C.

For cellulase and xylanase activities, plates containing 1.8% agar and 0.5% carboxymethyl cellulose (CMC) or 0.5% of xylan as substrate were prepared, respectively. These plates were subsequently stained for 20 min with 0.1% congo red under slight shaking and subsequently detained, first with 1 M NaCl for 20 min and then with 0.5% acetic acid, which was immediately rinsed off with cold water. Hydrolytic zones were observed against a red background.

MnP activity was determined in plates containing 1.8% agar, 20 mM 2,6-dimethoxyphenol (DMP), 5 mM MnSO_4_, and 10 mM H_2_O_2_ as substrates after the incubation at 30 °C. Laccase activity was determined similarly, but plates contained 1.8% agar and 1 mM guayacol as substrate. Oxidative dark brown zones were observed against the clear background as an indicator of laccase and peroxidase activity.

The secreted proteins were subjected to a native-PAGE (native, 10% acrylamide) for the detection of β-glucosidase activity. Electrophoresis was run at 12 mA and at 4 °C until the front dye left the gel. After running, the gel was soaked in 0.2 M sodium acetate buffer pH 5 for 10 min at room temperature. The gel was then incubated in 0.2 M sodium acetate buffer containing 0.1% (w/v) esculin and 0.03% (w/v) ferric chloride for 5 min at 50 °C. Alternatively, gels were incubated 10 min with sodium acetate buffer (0.2 M, pH 5) in the presence of either 1 or 2 M glucose, and then were revealed with the esculin solution keeping the same glucose concentration. Each reactive band was quantified by densitometric analysis using ImageJ software.

### 2.5 Secreted proteins extraction for proteomic analysis

Protein extraction was carried out according to Medina et al. (2004) with minor modifications. An inoculum of 1.10^6^ spores of *P. sanguineus* or *G. applanatum* was added to 20 mL of IM and incubated at 28 °C with rotary shaking (200 rpm) for 4 weeks. After this incubation period, the semi-solid medium obtained was extracted adding 5 mL of 50 mM sodium acetate buffer pH 4 containing 1 mM phenylmethylsulfonyl fluoride (PMSF) for 1 h at 4 °C under slight shaking. The secretome was recovered by filtration through filter paper and proteins were precipitated with 1 V of 20% trichloroacetic acid (TCA) and incubated at −20 °C ON. Subsequently, tubes were centrifuged at maximum speed for 10 minutes at 4 °C, washed three times with 70% ethanol, and finally with acetone, discarding the supernatant in each step. The pellet was dried at room temperature. Precipitated proteins were resuspended in 100 µL of 50 mM sodium acetate buffer pH 6 and were subjected to three sonication pulses of 5 seconds and 20% amplitude. Tubes were centrifuged at maximum speed and the supernatant was used for the next steps. First, a 20 µg aliquot of each sample was incubated for 45 min with 10 mM of DTT at 56 °C and then, with 20 mM of iodoacetamide at room temperature in darkness. Samples were sent to CEBIQUIEM (Facultad de Ciencias Exactas y Naturales, Universidad de Buenos Aires, Argentina) for further analysis.

### 2.6 LC-MS and data search analysis

The proteins were treated with trypsin and then cleaned with Zip-Tip C18 to extract the salts; the digested peptides were lyophilized by Speed Vac, and finally resuspended in 0.1% formic acid solution for mass spectrometric analysis. All solvents and reagents used were of LC-MS quality. The first peptide separation was carried out in a reverse phase capillary column (75 μm x 150 mm) packed with C18 (2,6 μm, 150 Å, Thermo Scientific). The flow rate was maintained at 300 nL/min. Mobile phase A (0.1% formic acid in water) and mobile phase B (0.1% formic acid in acetonitrile) were used to establish the 120 min gradient comprising 110 min of 5-35% B, 1 min 35-95% B, and 9 min of 95% B followed by re-equilibrating at 5% B for 5 min. The peptides were then analyzed by electrospray ionization (Easy-Spray, Thermo Scientific) with 3.5 kV potential. The peptides were separated in a second dimension by Mass Spectrometry (MS) (Q-Exactive, Thermo Scientific) with a High Collision Dissociation cell (HCD) and an Orbitrap analyzer. The equipment configuration allowed peptide identification simultaneously with chromatography resolution, obtaining Full MS and MS/MS. The spectra obtained were analyzed with “Proteome Discoverer” and database search was carried out in Uniprot (http://www.uniprot.org/) for *P. sanguineus* and against the functional annotation database of *G. lucidum* genome for *G. applanatum* (Chen et al., 2012). The precursor mass and MS/MS tolerances were 10 ppm and 0.05 Da, respectively. All the peptides were identified with a high confidence level. Two missed cleavage sites of trypsin were allowed. The oxidation (M) and carbamidomethylation (C) were set as variable and fixed modifications, respectively. The presence of a secretion signal peptide was determined with SignalP.

### 2.7 Statistical analysis

Data used for graphs of glucose released came from experiments repeated in triplicate and were analyzed using Sigma-Plot and GraphPad software. Inferential statistics was carried out applying a Two-way ANOVA test with a significance level < 0.1, to identify data with statistically significant differences.

## 3. Results and discussion

### 3.1 The Inductive Medium triggers a greater pretreatment efficiency using *P. sanguineus* secretome

Fungal secretomes from *G. applanatum* and *P. sanguineus* had previously been tested as biological pretreatment agents, resulting in considerable cellulose degradation levels in Argentinian rangeland species (Larran et al., 2015; Gauna et al., 2018). However, the fungal secretomes from those studies had been obtained in a culture medium with glucose as a carbon source - Potato Glucose (PG) medium-, which is known to transcriptionally repress cellulases, hemicellulases, and other lignocellulolytic enzymes in filamentous fungi (Suzuki et al., 2008; Glass et al., 2013). Moreover, according to previous studies, the absence of glucose and the presence of a lignocellulosic substrate may induce the expression of lignocellulolytic enzymes (Suzuki et al., 2008; Rohr et al., 2013).

In this work, it has been developed and tested an inductive medium (IM) with *P. prionitis* leaves as a lignocellulosic substrate, which notoriously increased glucose release levels in comparison with PG medium. The results show that glucose release levels depend on the fungal secretome and the biomass used, reaching the highest values with *P. sanguineus* secretome acting on green leaves (Fig. 1). The greater amount of lignin in senescent leaves could hinder accessibility to polysaccharides, causing decay in glucose release. Differences in glucose released between fungal secretomes could be due to the higher protein concentration in *P. sanguineus* secretome (see section 3.4), although some differential enzymatic activities could also affect the total process, as discussed next.

**Figure 1.**
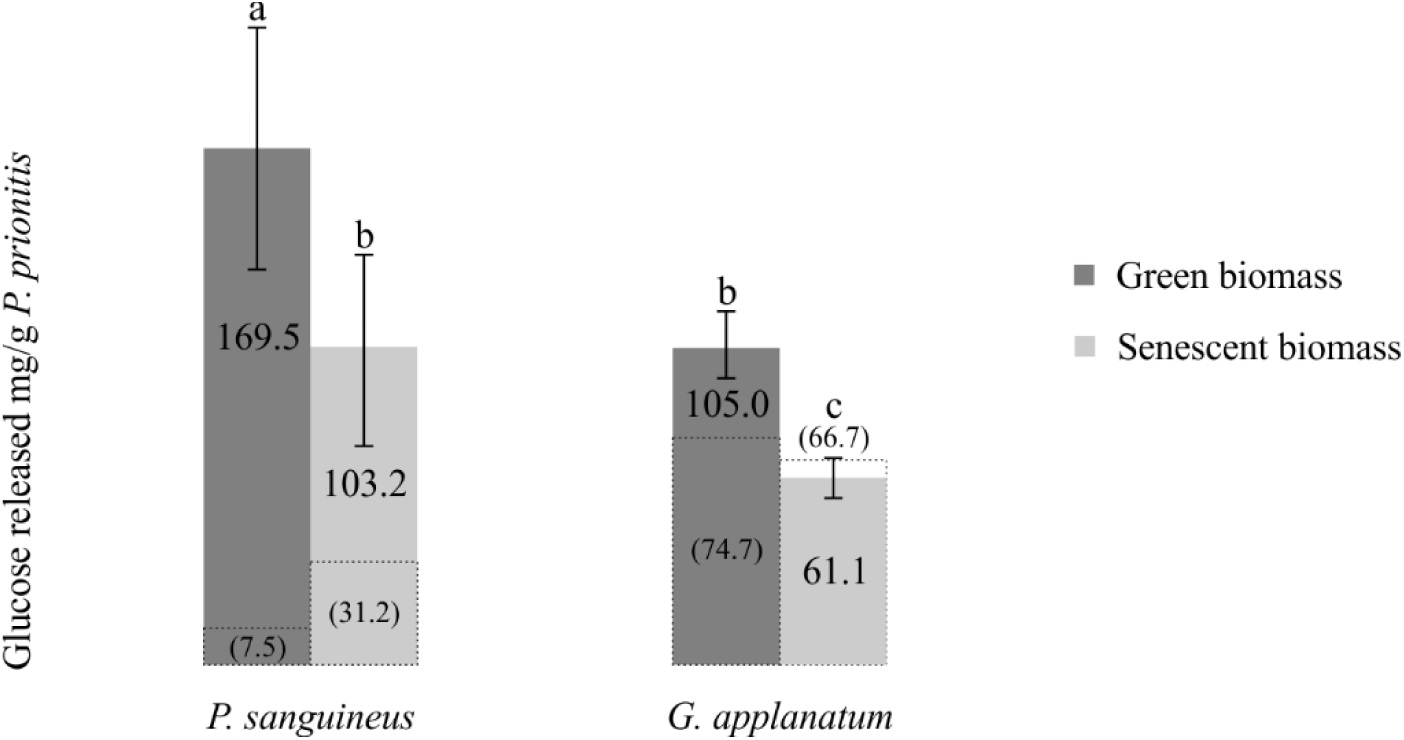
Total glucose released with fungal secretomes obtained in IM. Numbers in brackets indicate the total glucose released with fungal secretomes obtained in PG medium (Gauna et al., 2018). Error bars represent the standard error. Different letters indicate statistically significant differences.

The use of *P. sanguineus* secretome obtained in IM allowed a 22.5-fold and a 3-fold increase in glucose release compared to the secretome produced in PG medium over green and senescent biomass, respectively (Gauna et al., 2018) (Fig. 1). Regarding *G. applanatum* secretome, the increase in glucose released respect to PG medium was only 1.4-fold for green biomass and no significant changes were observed for senescent biomass. Differences in the total glucose released between the two culture media, PG vs IM, may be explained by the catabolic repression process as well as the inducible lignocellulolytic enzymes, as described above.

The comparison between the percent of hydrolyzed cellulose in IM vs. PG medium allows an easier appreciation of the differential behavior of each fungus (Table 1). The contribution of *P. sanguineus* secretome to the pretreatment step was notoriously higher than the *G. applanatum* secretome contribution, similarly to the behavior observed in previous studies (Gauna et al., 2018). The higher contribution of *P. sanguineus* secretome to the pretreatment process could be due to an increased production/activity of lignin degrading enzymes; whereas in the case of *G. applanatum*, the higher contribution to the hydrolysis step could be due to a proportionally greater cellulolytic activity. The IM tested in this work triggers an increase in the pretreatment contribution of *P. sanguineus* secretome respect to PG medium (up to 80 and 85 % vs. 34 and 59 % in green and senescent biomass, respectively). This shows that *P. sanguineus* enzyme consortium involved in lignin degradation is preferentially inducible by the presence of a lignocellulosic substrate and the absence of glucose. In contrast, no differences in pretreatment contribution were observed with *G. applanatum* secretome between both media.

**Table 1.**
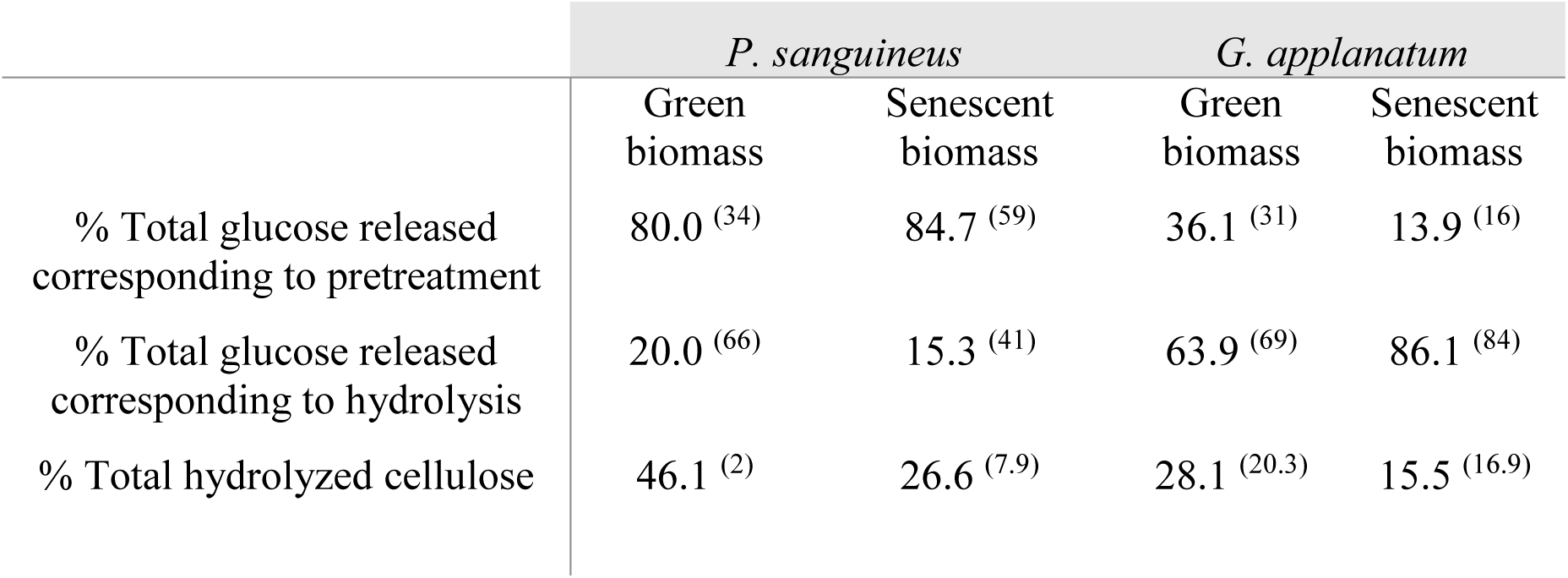
Contribution of pretreatment and hydrolysis steps to glucose release on green and senescent *P. prionitis* biomass. The numbers in brackets correspond to the values obtained in PG medium previously (Gauna et al., 2018).

### 3.2 The activity of lignocellulolytic enzymes is higher in *P. sanguineus* secretome

The activity of the main lignocellulolytic enzymes identified in the secretomes was assessed in Petri dishes. The employment of wheat bran (WB) as a carbon source was previously evaluated and proved to substantially contribute to the expression of lignocellulolytic enzymes (Scarpa et al., 2019). Therefore, WB was used as a control to evaluate the different activities from IM. Figure 2 shows the plate assays for cellulase, xylanase, laccase, and MnP activities. Only the cellulase activity was observed in both secretomes with both carbon sources, with the greatest activity corresponding to *P. sanguineus* secretome. Curiously, xylanase activity was detected in both secretomes from fungi growing in IM, while a very low activity was observed in WB only in *P. sanguineus* secretome. Laccase and MnP activities were only detected in *P. sanguineus* secretome with both carbon sources, being greater in WB medium.

**Figure 2.**
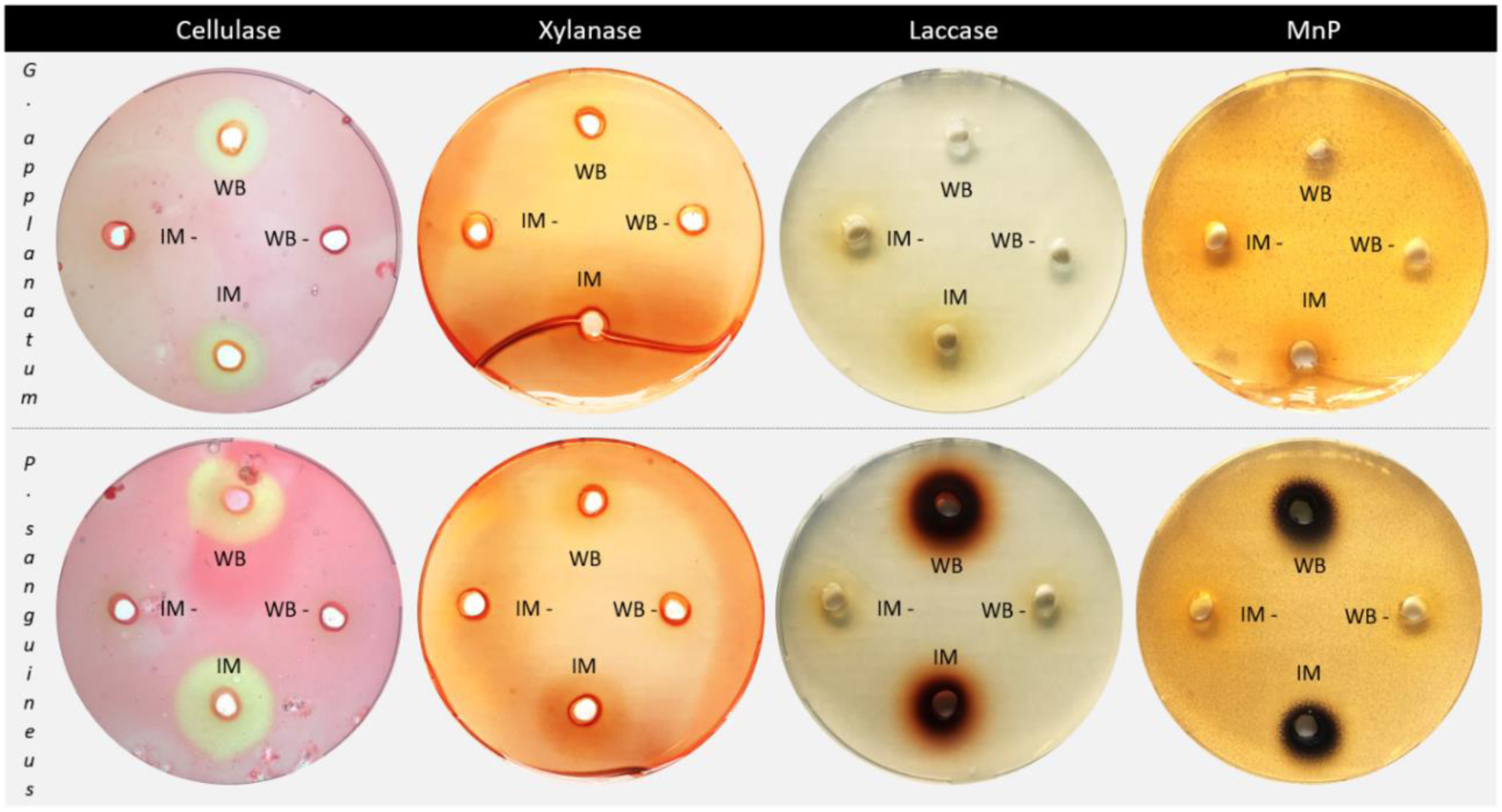
Plate assays of lignocellulosic activities. WB and IM: different culture media and their negative controls (WB-, IM-). Twenty μg of total secreted proteins were added into different wells within each plate.

The exclusive detection of MnP and laccase activities in *P. sanguineus* secretome could provide a rational explanation for its greater contribution to the pretreatment process compared to *G. applanatum* secretome, as stated in section 3.1. In compliance with the lower percentage of hydrolyzed cellulose, *G. applanatum* showed a reduced activity of all the enzymes assayed regarding *P. sanguineus*.

### 3.3 *P. sanguineus* secretes a putatively glucose-tolerant β-glucosidase

β-glucosidases play a critical role among the enzymes involved in plant biomass deconstruction, since they catalyze the last step of sugar degradation before the fermentation phase (Santos et al., 2019), mitigating product inhibition of cellulases and hemicellulases. However, most fungal β-glucosidases characterized show a weak activity at high glucose concentrations, limiting their enzymatic hydrolysis capacity in industrial processes (Xiao et al., 2004). Besides, fungi do not generally secrete large quantities of β-glucosidases into the extracellular medium (Cairns and Esen, 2010; Jeya et al., 2010). Hence, numerous studies have focused on the differential characteristics (including sequence and structural data) that would lead to glucose tolerance, in order to find better natural producers and/or improve the engineering strategies of these enzymes (de Giuseppe et al., 2014; Guo et al., 2016; Mariano et al., 2017; Salgado et al., 2018; Santos et al., 2019).

To investigate the presence of any glucose-tolerant β-glucosidase isoform, it has been performed zymograms for this activity under different glucose concentrations. As shown in Figure 3, the highest β-glucosidase activity was achieved with WB as a carbon source on *P. sanguineus* secretome, like laccase and MnP activities (see section 3.2). This secretome shows two isoforms, the high molecular weight (HMW) isoform presented the highest β-glucosidase activity in almost all the conditions. The exception was WB medium at glucose 2M, where the contribution of each isoform was equivalent (50%, lane 11 from Fig. 3A).

**Figure 3.**
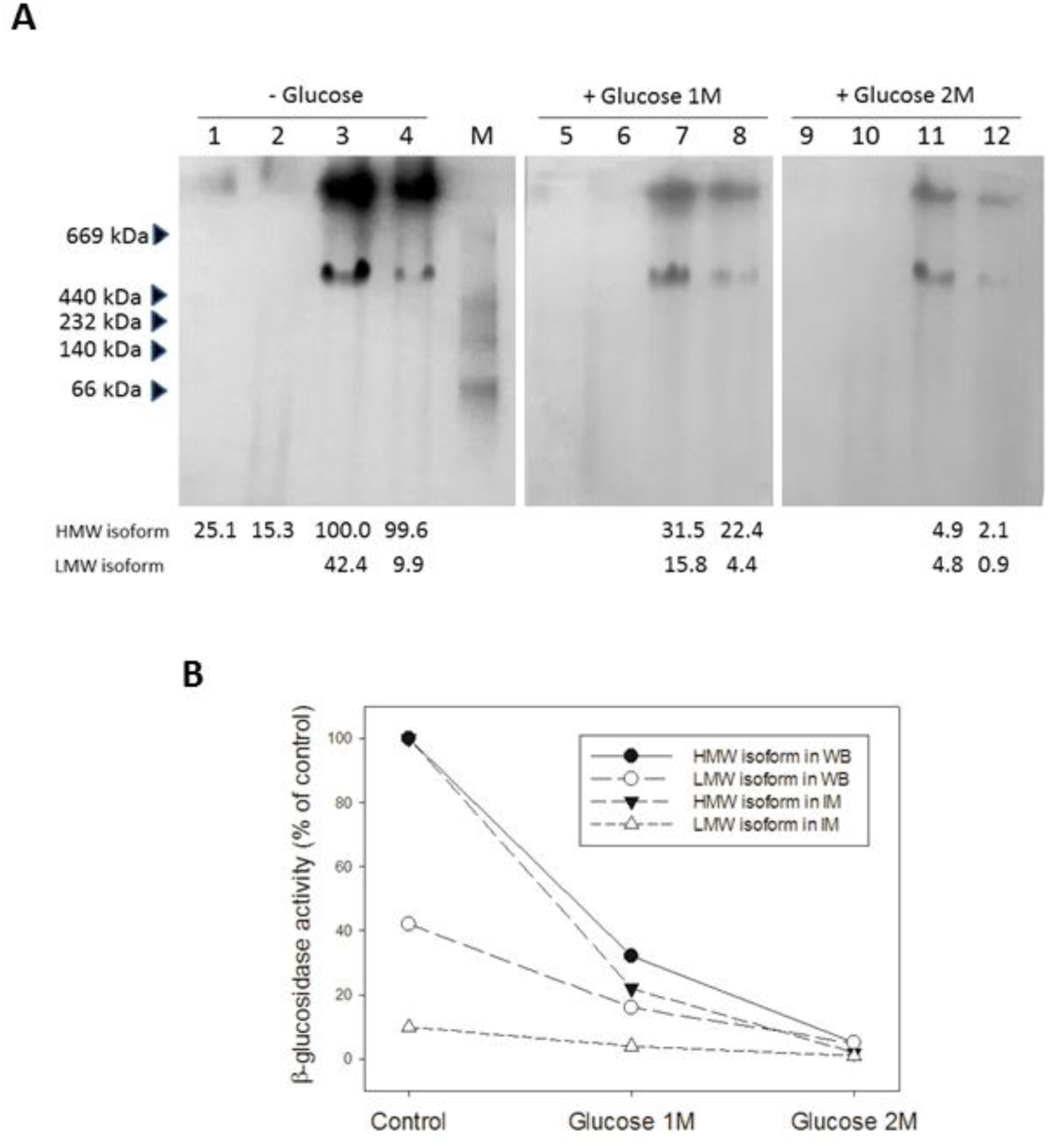
**A)** Non-denaturing polyacrylamide gel electrophoresis stained by β-glucosidase activity in absence/presence of glucose. Twenty μg of total secreted proteins were added into different lanes. *G. applanatum* secretomes were evaluated in lanes 1, 2, 5, 6, 9, and 10; while *P. sanguineus* secretomes were evaluated in lanes 3, 4, 7, 8, 11, and 12. Odd number lanes were loaded with secretomes from WB medium; while even number lanes contained secretomes from IM. Quantitative zymogram analysis for each isoform relative to HMW isoform from lane 3 is shown at the bottom. M: molecular weight markers **B)** Decreasing β-glucosidase activity from each isoform at increasing glucose concentrations.

The residual activity of each isoform at increasing glucose concentrations showed that the HMW isoform is notoriously affected by this metabolite (2-5 % residual activity at glucose 2M). In contrast, the low molecular weight (LMW) isoform showed a less steep decrease in both tested media (Fig. 3B), suggesting to be a glucose-tolerant β-glucosidase.

This finding opens new avenues for second-generation biofuel production regarding the usual limitations of product inhibition for cellulases and hemicellulases. Curiously, this isoform identified as a GH3 family member by MALDI-TOF analysis (Table S1), showed a 10% residual activity at very high glucose concentrations (2M). Although most glucose-tolerant β-glucosidases characterized belong to the GH1 family (Mariano et al., 2017; Salgado et al., 2018), several exceptions in which they are classified as GH3 family members were reported (Huang et al., 2014; Valappil et al., 2019; Monteiro et al., 2020).

Moreover, in agreement with the presented results, a recent report has characterized a secreted β-glucosidase from *Aspergillus unguis*, identified as a GH3 that resulted in a LMW isoform with intact activity even at 0.5 M glucose. Also, docking analysis supported its high glucose tolerance (Valappil et al., 2019). However, future studies are necessary to assess the significance of these findings.

On the other hand, *G. applanatum* only exhibited the HMW β-glucosidase isoform, which showed notoriously lower activity levels than those of *P. sanguineus*. This result could be related either to a lower amount of secreted protein and/or a lower specific activity. In any case, the low basal activity made it impossible to detect this isoform in the presence of glucose.

### 3.4 Proteomic characterization reveals many functional biomass-degrading enzymes and uncharacterized proteins in both fungal secretomes

LC-MS analysis allowed the identification of 180 proteins ranging from 12.2 to 234.3 kDa and 73 proteins from 13.6 to 223.8 kDa in *P. sanguineus* and *G. applanatum* secretomes, respectively. Calculated pI varied from 4 to 8. In both cases, proteins were obtained from IM, and protein concentration in *P. sanguineus* secretome resulted three-fold higher than in *G. applanatum* secretome (1.07 mg.mL^-1^ vs. 0.31 mg.mL^-1^, respectively). Interestingly, a simulated 2D gel presentation of each secretome from the LC-MS data fitted the 2D gels obtained experimentally (Fig. S1), validating the coverage of the method used.

The proteomic approach, carried out for the first time in *P. sanguineus* and *G. applanatum*, allowed a deep characterization of the secretomes from these filamentous fungi. Proteins identified in each secretome and their corresponding peptide sequences are fully listed in supplementary tables S2 and S3. The majority presented a signal peptide. Proteins identified were functionally classified according to their biological role. Figure 4 depicts the functional classification of *P. sanguineus* and *G. applanatum* secreted proteins, which were grouped into cellulose, hemicellulose or lignin-degrading enzymes, glycoside hydrolases, uncharacterized proteins, and others (miscellaneous). It is worth remarking the significant proportion of uncharacterized proteins found in both species. Tables 2 and 3 summarize the characterized proteins from *P. sanguineus* and *G. applanatum* secretomes, respectively.

**Table 2.**
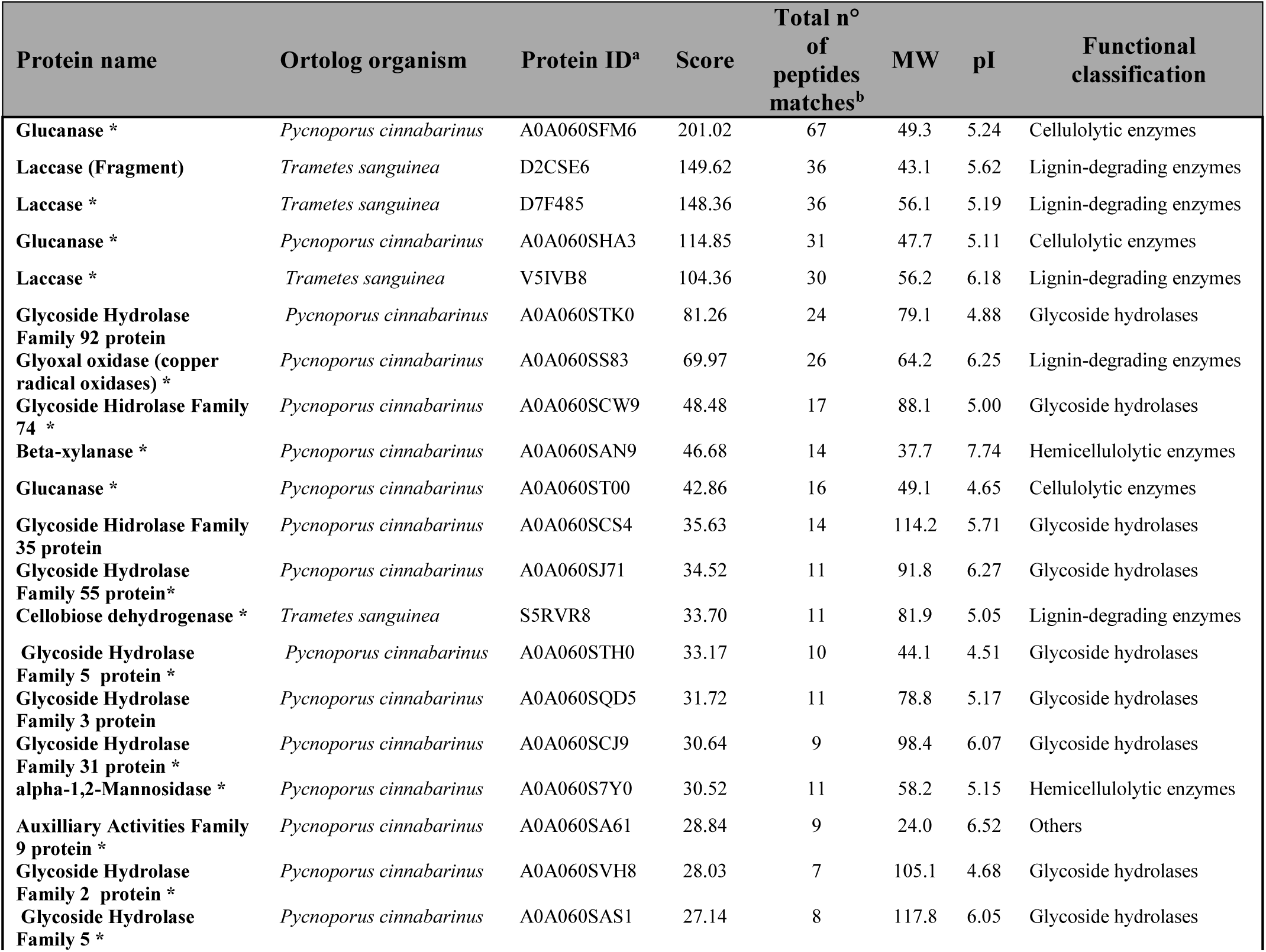
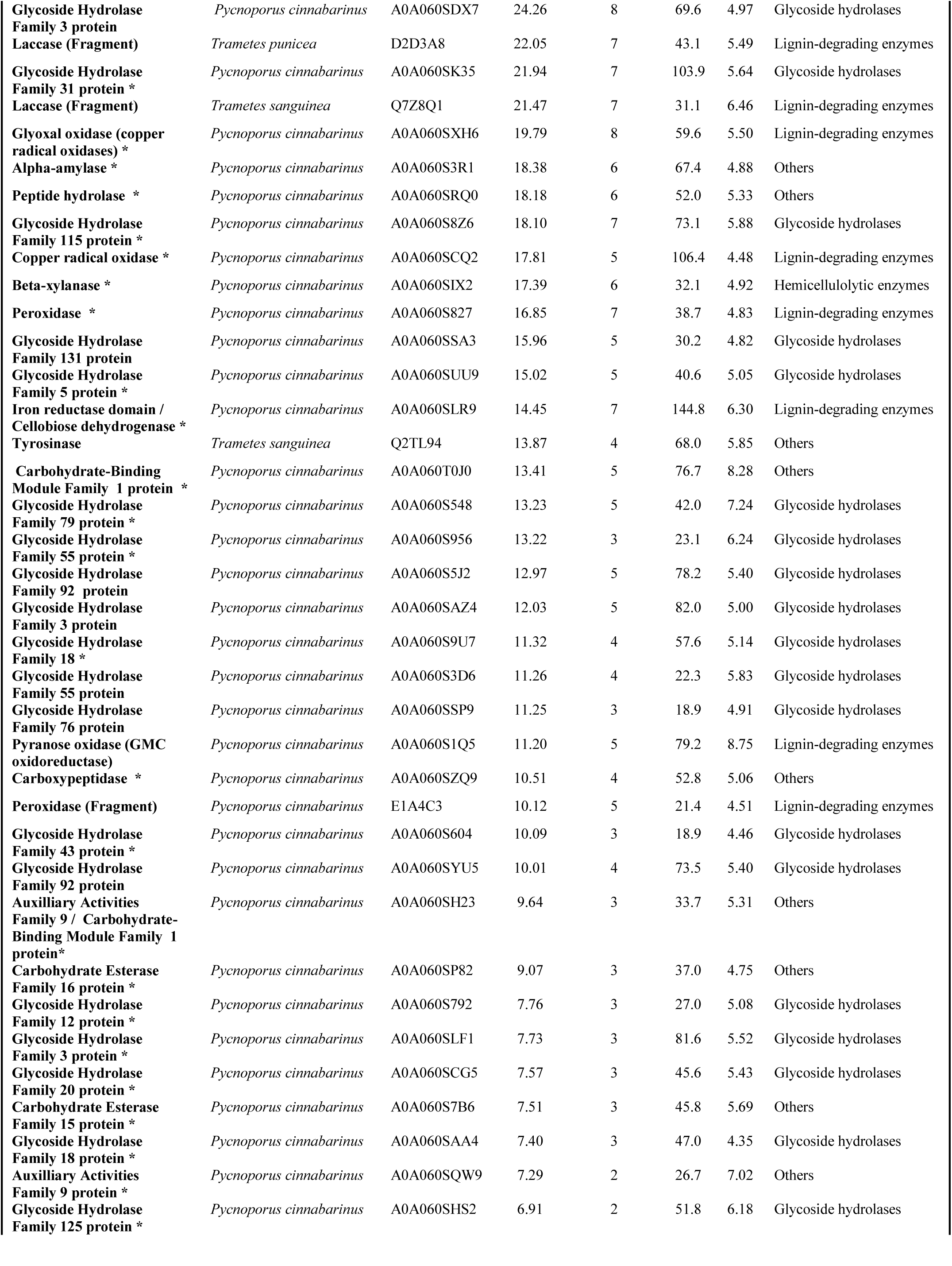
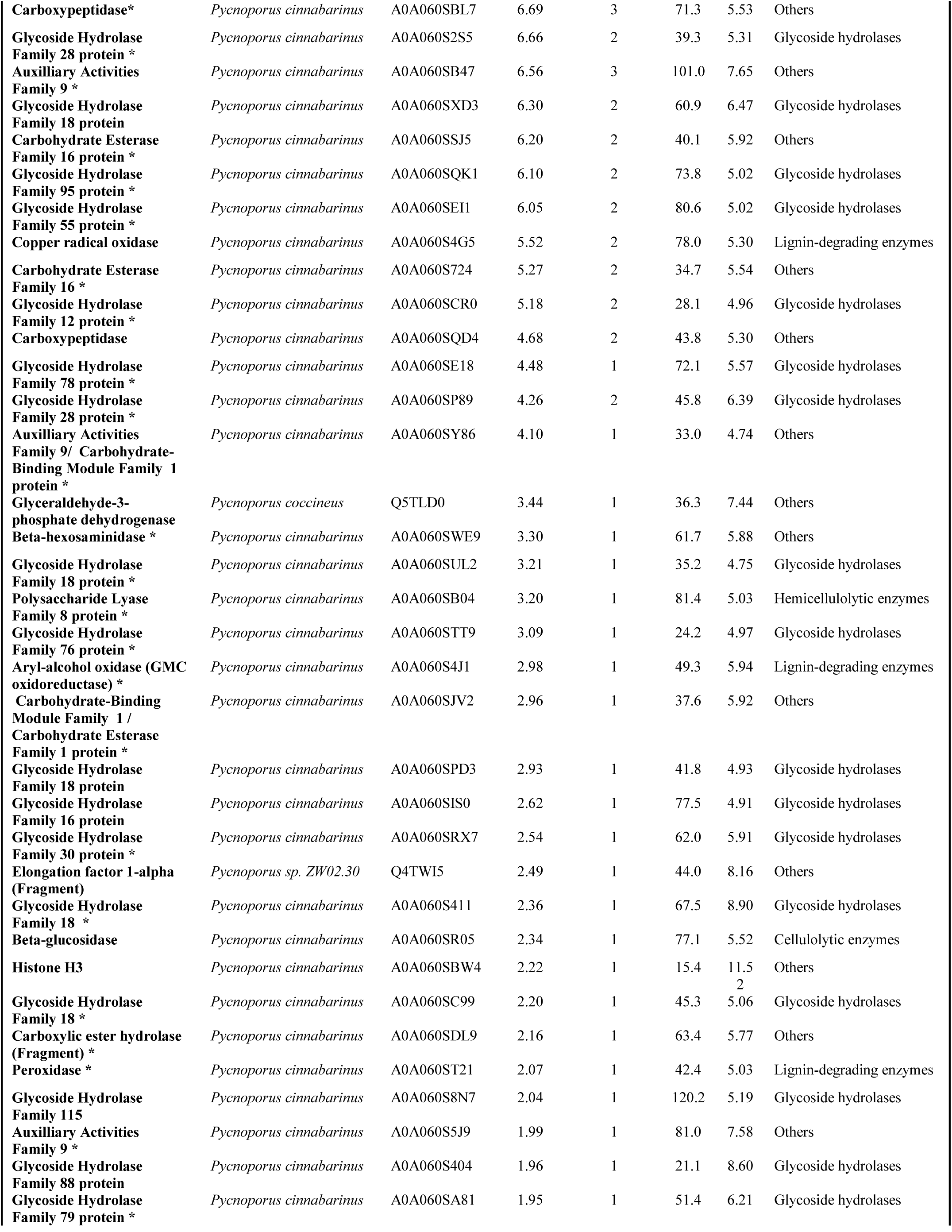
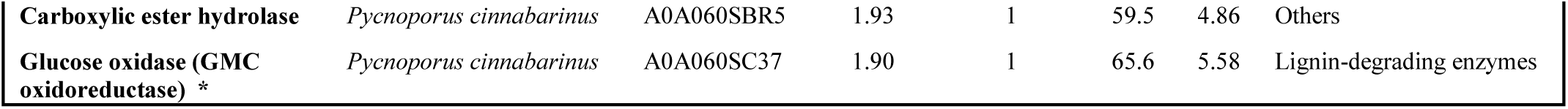
Detailed information of 94 characterized proteins from *P. sanguineus* secretome. ^a^ matched protein ID was derived from the Uniprot database. ^b^ the sequences of matched peptides are shown in Table S2. ^*^ indicates the presence of a signal peptide in the matched protein

**Table 3.**
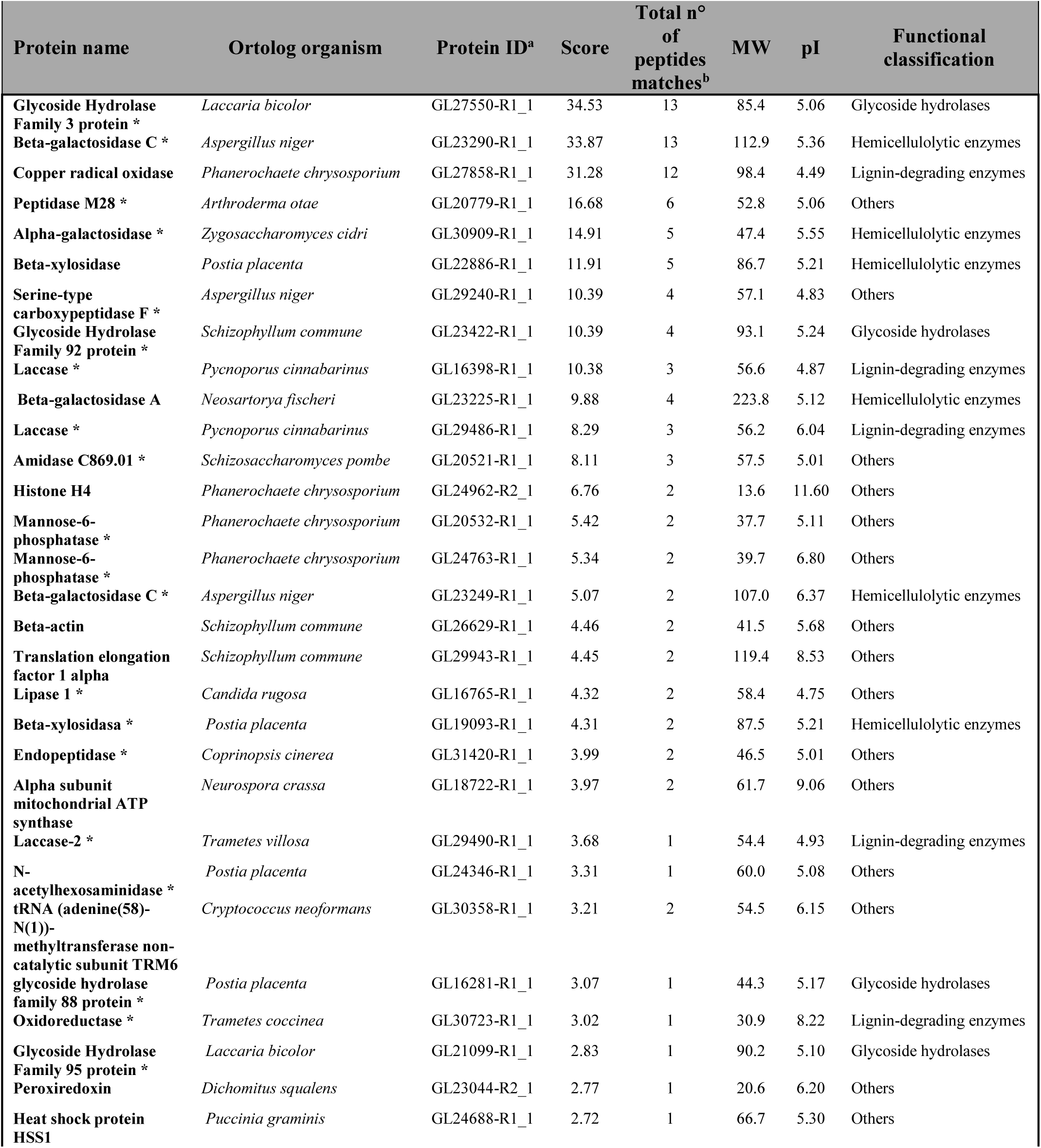
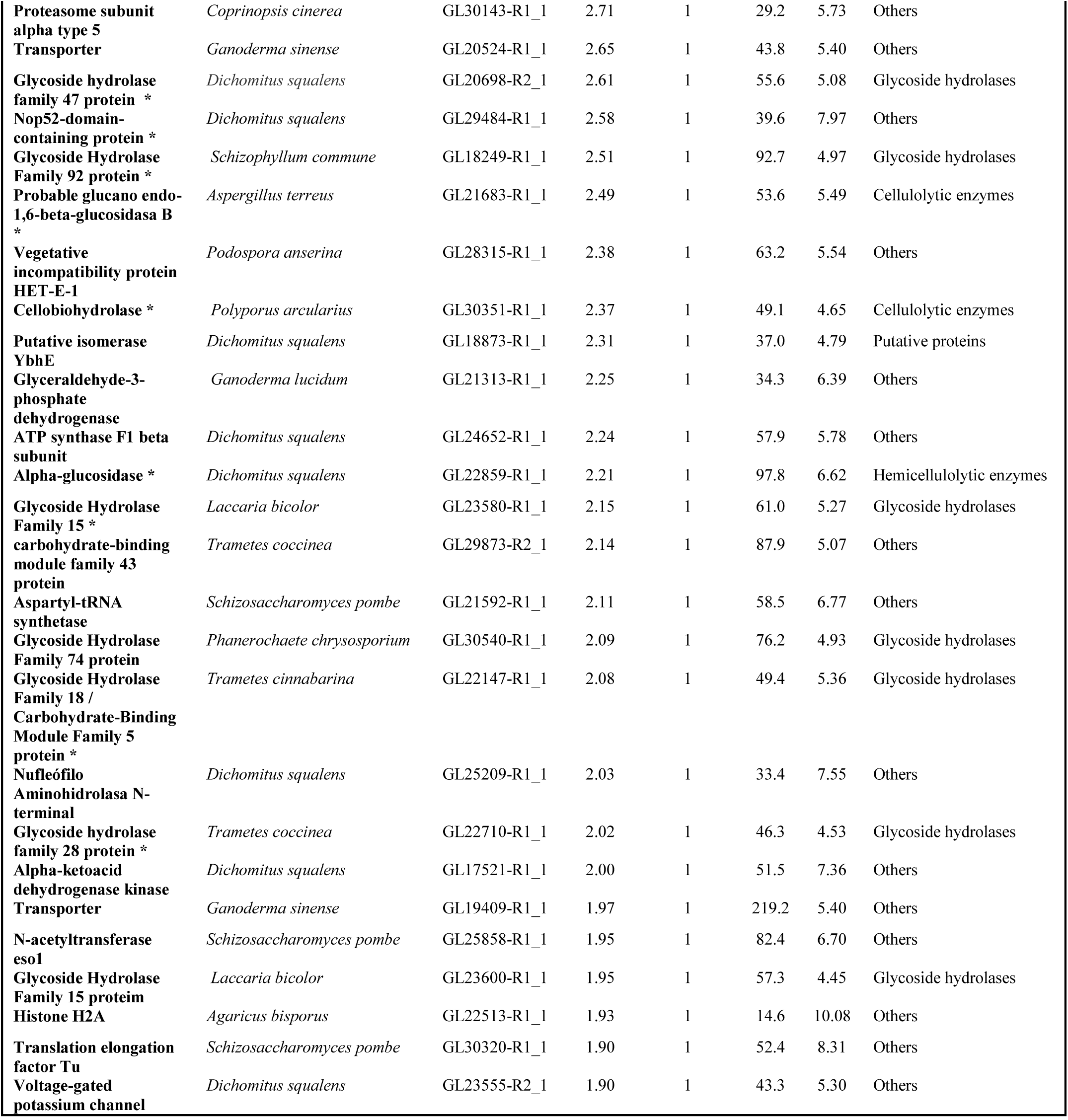
Detailed information of 56 characterized proteins from G. *applanatum* secretome. ^a^ matched protein ID was derived from the *G. lucidum* genome database (Chen et al. 2012). ^b^ the sequences of matched peptides are shown in Table S3. ^*^ indicates the presence of a signal peptide in the matched protein

**Figure 4.**
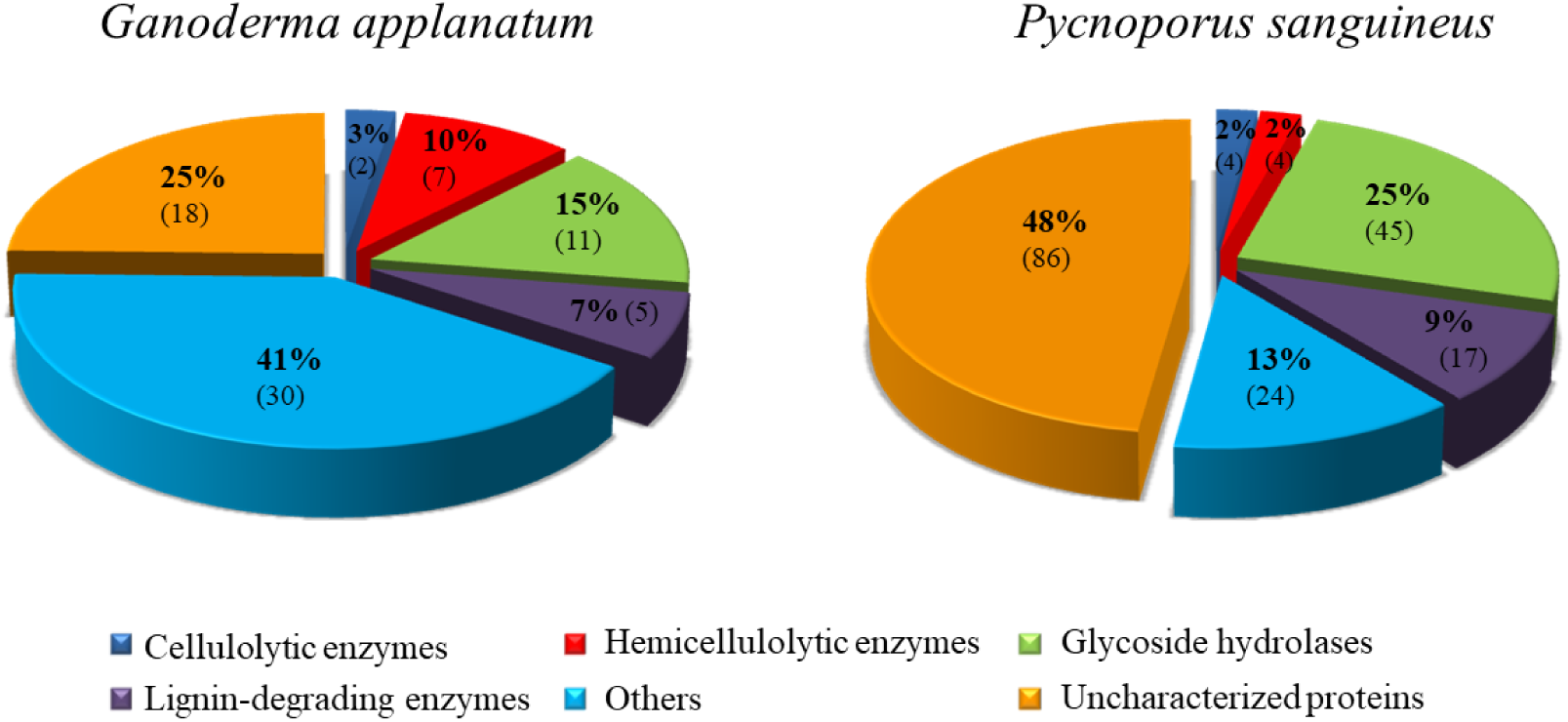
Functional classification of the secreted proteins according to their biological role. The numbers in brackets correspond to the number of proteins identified in each group.

Glycoside hydrolases (GH) are a widespread group of enzymes that catalyze the hydrolysis of O- or S-glycosides. They can be classified according to different criteria: sequence or structure-based methods (most used), the stereochemical outcome of the hydrolysis reaction, and exo or endo acting (Cantarel et al., 2009). Given the broad substrate specificity of these enzymes, they can act in several metabolic pathways, such as cellulose, hemicellulose, and lignin degradation. Coherently, they represented a meaningful category in both secretomes (15-25% of total proteins).

Cellulases include three major groups of enzymes: endo-glucanases, exo-glucanases (or cellobiohydrolases), and β-glucosidases, all involved in the conversion of cellulose into glucose (Behera et al., 2017). As was discussed previously, in wood-degrading basidiomycetes, cellulases expression is regulated by carbon catabolite repression mechanisms and by the presence of lignocellulosic substrates (Rohr et al., 2013). Two enzymes from this family (an endo-1,6-β-glycosidase and a cellobiohydrolase) were found in the secretome of *G. applanatum*, while four (three glucanases and a β-glycosidase) were identified in the *P. sanguineus* secretome.

Complete hydrolysis of hemicellulose requires of several enzymes: endo-1,4-β-xylanases, exo-1,4-β-xylanases, β-xylosidases, β-arabynofuranosidases, β-d-gucorinidases, acetylxylanoesterases, galactosidases, mannanases, mannosidases, polysaccharide lyases, phytases, among others (Couturier and Berrin, 2013). Similarly to cellulases, hemicellulases require the presence of a lignocellulosic compound to be transcriptionally induced. At least seven proteins were linked to hemicellulose hydrolysis in the *G. applanatum* secretome. It has been found three β-galactosidases, two β-xylosidases, an α-galactosidase, and an α-glucosidase. On the other hand, two xylanases, a β-1,2-mannosidase, and a polysaccharide lyase were detected in the *P. sanguineus* secretome. In brief, at least 10 and 2% of the different proteins identified in *G. applanatum* and *P. sanguineus* secretomes, respectively, belong to the hemicellulose-degrading enzymes category. However, these percentages are actually higher because the GH identified in both secretomes would be mainly involved in this process, as shown by the most represented families (GH3, GH18, GH55, GH92) (http://www.cazy.org, Jul 2020).

Several enzymes are involved in lignin breakdown, such as laccases, peroxidases, glycose oxidases, isoamyl-alcohol-oxidases, glutathione reductases, and glutathione-S-transferases. Lacasse, lignin peroxidase, and MnP are considered the most powerful lignin-degrading enzymes (Dashtban et al., 2010). In this study, it has been identified three laccases, a copper radical oxidase, and a non-characterized oxidoreductase in *G. applanatum* secretome. Regarding *P. sanguineus* secretome, five laccases, two cellobiose dehydrogenases, three peroxidases, three glycose-methanol-coline (GMC) oxidoreductases, and four copper radical oxidases were found.

The steps involved in hydrogen peroxide production are particularly important for lignin degradation because this substrate is needed for the catalytic activity of peroxidases. GMC oxidoreductases and copper radical oxidases are involved in such production. Interestingly, a non-characterized GMC oxidoreductase in *G. applanatum* secretome, as well as a glucose oxidase, an aryl-alcohol-oxidase, and a pyranose oxidase were identified in *P. sanguineus* secretome. As already mentioned, four copper radical oxidases were also found in *P. sanguineus* secretome, while two of them were glyoxal oxidases. The cellobiose dehydrogenase (CDH) detected in *P. sanguineus* secretome is an extracellular redox enzyme that plays a major role in lignin degradation by breaking beta-ethers, demethoxylating aromatic structures in lignin, and introducing hydroxyl groups in non-phenolic lignin. Moreover, CDH generates hydroxyls by reducing Fe^3+^ to Fe^2+^ and O_2_ to H_2_O_2_ and may be converted into a quinone-oxidoreductase by proteolysis (Moukha et al., 1999; Manavalan et al., 2012).

Consistently, some proteases were identified (peptidases, endopeptidases, and carboxypeptidases), which were previously associated with lignocellulose degradation (Dosoretz et al., 1990) by activating cellulases and cleaving functional domains of the CDH (Habu et al., 1993; Manavalan et al., 2012).

In addition to all these enzymes, many uncharacterized proteins with unknown function were found in both species, especially in *P. sanguineus* secretome (48% of the total). The substantial percentage of uncharacterized proteins indicates the low degree of knowledge of these species, reflecting that a greater amount and depth of experimental work in the subject is needed for a better comprehension and application of the mechanisms underlying lignocellulosic biomass degradation.

## 4. Conclusions

In this study, *Pycnoporus sanguineus* and *Ganoderma applanatum* secretomes were evaluated as pretreatment agents in a fermentable-sugar obtaining process. The inductive medium containing *Panicum prionitis* leaves increased the percentage of hydrolyzed cellulose compared with a non-inductive medium. The pretreatment efficiency and the activities of the main hydrolytic enzymes were higher using *P. sanguineus* secretome. Diverse proteins were identified and functionally classified according to their biological role. Many uncharacterized proteins were found, reflecting the still insufficient knowledge of the mechanisms underlying lignocellulose degradation. Interestingly, a putative glucose-tolerant β-glucosidase was identified in *P. sanguineus* secretome, opening new avenues for second-generation biofuel production.

## Acknowledgements

This work was supported by AGENCIA NACIONAL DE PROMOCION CIENTIFICA Y TECNOLOGICA (D-TEC 2013 N° 0001/13) and SECTEI-Santa Fe Province (Res 156/2017). The funding source had no involvement in study design. Authors would like to thank their supporting organisms and institutions. AG is a fellow of Fundación Ciencias Agrarias. ASL is a fellow of Consejo Nacional de Investigaciones Científicas y Técnicas (CONICET). VEP is a Professor at Universidad Nacional de Rosario (UNR). SRF and HRP are Professors at Universidad Nacional de Rosario (UNR) and researchers of IICAR-CONICET.

