## Supplementary material for "Secretome characterization of the lignocellulose-degrading fungi *Pycnoporus sanguineus* and *Ganoderma applanatum*": Table S1

| Band | Protein identify | *Ortolog organism* | Protein ID | Score | Total n° of peptides matches | MW | pI | Estimated Native MW[a] |
|---|---|---|---|---|---|---|---|---|
| **HMW** | **Glycoside hydrolase family 3 protein** | *Pycnoporus cinnabarinus* | A0A060SDX7 | 2.12 | 1 | 69.6 | 4.97 | >669 kDa (octamer?) |
| **LMW** | **Beta-glucosidase** | *Pycnoporus cinnabarinus* | A0A060SR05 | 2.08 | 1 | 77.1 | 5.52 | >440 kDa (hexamer?) |

Table S1. Protein identification of the two β-glucosidase activity bands from *P. sanguineus* secretome. Detailed data for these proteins can be found in Table S2.

[a] Note that it is an speculation based on the electrophoretic mobility, but it is necessary an exclusion chromatography to determine this parameter.
