## Supplementary material for "Secretome characterization of the lignocellulose-degrading fungi *Pycnoporus sanguineus* and *Ganoderma applanatum*": Table S2

Table S2. Detailed information of 180 proteins identified from P. *sanguineus* secretome

| Accession | Description | Score | Coverage | # Proteins | # Unique Peptides | # Peptides | # PSMs | # AAs | MW [kDa] | calc. pI |
|---|---|---|---|---|---|---|---|---|---|---|
| A0A060SFM6 | Glucanase OS=Pycnoporus cinnabarinus GN=BN946_scf185007.g75 PE=3 SV=1 - [A0A060SFM6_PYCCI] | 201,02 | 37,06 | 2 | 10 | 13 | 67 | 456 | 49,3 | 5,24 |
| D2CSE6 | Laccase (Fragment) OS=Trametes sanguinea PE=4 SV=1 - [D2CSE6_9APHY] | 149,62 | 35,53 | 28 | 1 | 9 | 36 | 394 | 43,1 | 5,62 |
| D7F485 | Laccase OS=Trametes sanguinea PE=3 SV=1 - [D7F485_9APHY] | 148,36 | 26,64 | 18 | 1 | 8 | 36 | 518 | 56,1 | 5,19 |
| A0A060SHA3 | Glucanase OS=Pycnoporus cinnabarinus GN=BN946_scf185014.g56 PE=3 SV=1 - [A0A060SHA3_PYCCI] | 114,85 | 25,83 | 1 | 11 | 11 | 31 | 453 | 47,7 | 5,11 |
| V5IVB8 | Laccase OS=Trametes sanguinea GN=lcc1 PE=2 SV=1 - [V5IVB8_9APHY] | 104,36 | 39,96 | 5 | 8 | 11 | 30 | 518 | 56,2 | 6,18 |
| A0A060SE04 | Uncharacterized protein OS=Pycnoporus cinnabarinus GN=BN946_scf184748.g21 PE=4 SV=1 - [A0A060SE04_PYCCI] | 87,55 | 11,91 | 1 | 4 | 4 | 25 | 571 | 61,7 | 5,41 |
| A0A060STK0 | Glycoside Hydrolase Family 92 protein OS=Pycnoporus cinnabarinus GN=BN946_scf184969.g71 PE=4 SV=1 - [A0A060STK0_PYCCI] | 81,26 | 22,65 | 1 | 9 | 9 | 24 | 724 | 79,1 | 4,88 |
| A0A060SS83 | Glyoxal oxidase OS=Pycnoporus cinnabarinus GN=BN946_scf184747.g48 PE=4 SV=1 - [A0A060SS83_PYCCI] | 69,97 | 16,58 | 2 | 6 | 9 | 26 | 597 | 64,2 | 6,25 |
| A0A060SEX1 | Uncharacterized protein (Fragment) OS=Pycnoporus cinnabarinus GN=BN946_scf184829.g57 PE=3 SV=1 - [A0A060SEX1_PYCCI] | 58,18 | 25,35 | 1 | 6 | 6 | 18 | 505 | 56,0 | 5,29 |
| A0A060SCW9 | Glycoside Hydrolase Family 74 / Carbohydrate-Binding Module Family 1 protein OS=Pycnoporus cinnabarinus GN=BN946_scf184977.g4 PE=4 SV=1 - [A0A060SCW9_PYCCI] | 48,48 | 18,20 | 1 | 11 | 11 | 17 | 846 | 88,1 | 5,00 |
| A0A060SAN9 | Beta-xylanase OS=Pycnoporus cinnabarinus GN=BN946_scf184817.g18 PE=3 SV=1 - [A0A060SAN9_PYCCI] | 46,68 | 8,33 | 1 | 2 | 2 | 14 | 360 | 37,7 | 7,74 |
| A0A060ST00 | Glucanase OS=Pycnoporus cinnabarinus GN=BN946_scf184969.g23 PE=3 SV=1 - [A0A060ST00_PYCCI] | 42,86 | 12,23 | 2 | 2 | 5 | 16 | 458 | 49,1 | 4,65 |
| A0A060SCS4 | Glycoside Hydrolase Family 35 protein OS=Pycnoporus cinnabarinus GN=BN946_scf184970.g129 PE=3 SV=1 - [A0A060SCS4_PYCCI] | 35,63 | 9,30 | 1 | 8 | 8 | 14 | 1054 | 114,2 | 5,71 |
| A0A060ST91 | Uncharacterized protein OS=Pycnoporus cinnabarinus GN=BN946_scf184775.g22 PE=3 SV=1 - [A0A060ST91_PYCCI] | 34,99 | 11,70 | 2 | 3 | 3 | 11 | 393 | 41,6 | 4,81 |

| | | | | | | | | | | |
|---|---|---|---|---|---|---|---|---|---|---|
| A0A060SJ71 | Glycoside Hydrolase Family 55 protein OS=Pycnoporus cinnabarinus GN=BN946_scf184980.g29 PE=4 SV=1 - [A0A060SJ71_PYCCI] | 34,52 | 7,28 | 1 | 5 | 5 | 11 | 865 | 91,8 | 6,27 |
| S5RVR8 | Cellobiose dehydrogenase OS=Trametes sanguinea PE=2 SV=1 - [S5RVR8_9APHY] | 33,70 | 11,70 | 3 | 4 | 8 | 11 | 769 | 81,9 | 5,05 |
| A0A060STH0 | Carbohydrate-Binding Module Family 1 / Glycoside Hydrolase Family 5 protein OS=Pycnoporus cinnabarinus GN=BN946_scf184921.g29 PE=3 SV=1 - [A0A060STH0_PYCCI] | 33,17 | 12,17 | 1 | 3 | 3 | 10 | 419 | 44,1 | 4,51 |
| A0A060SQD5 | Glycoside Hydrolase Family 3 protein OS=Pycnoporus cinnabarinus GN=BN946_scf184868.g9 PE=4 SV=1 - [A0A060SQD5_PYCCI] | 31,72 | 9,15 | 1 | 5 | 5 | 11 | 732 | 78,8 | 5,17 |
| A0A060SCJ9 | Glycoside Hydrolase Family 31 protein OS=Pycnoporus cinnabarinus GN=BN946_scf184940.g83 PE=3 SV=1 - [A0A060SCJ9_PYCCI] | 30,64 | 8,52 | 1 | 4 | 4 | 9 | 892 | 98,4 | 6,07 |
| A0A060S7Y0 | alpha-1,2-Mannosidase OS=Pycnoporus cinnabarinus GN=BN946_scf184569.g13 PE=3 SV=1 - [A0A060S7Y0_PYCCI] | 30,52 | 21,28 | 1 | 7 | 7 | 11 | 531 | 58,2 | 5,15 |
| A0A060SF78 | Uncharacterized protein OS=Pycnoporus cinnabarinus GN=BN946_scf185002.g21 PE=4 SV=1 - [A0A060SF78_PYCCI] | 30,45 | 8,47 | 1 | 4 | 4 | 8 | 602 | 64,9 | 5,31 |
| A0A060SA61 | Auxilliary Activities Family 9 protein OS=Pycnoporus cinnabarinus GN=BN946_scf184908.g125 PE=4 SV=1 - [A0A060SA61_PYCCI] | 28,84 | 8,70 | 1 | 3 | 3 | 9 | 230 | 24,0 | 6,52 |
| A0A060SVH8 | Glycoside Hydrolase Family 2 protein OS=Pycnoporus cinnabarinus GN=BN946_scf184615.g9 PE=4 SV=1 - [A0A060SVH8_PYCCI] | 28,03 | 5,53 | 1 | 3 | 3 | 7 | 958 | 105,1 | 4,68 |
| A0A060SAS1 | Carbohydrate-Binding Module Family 1 / Glycoside Hydrolase Family 5 protein OS=Pycnoporus cinnabarinus GN=BN946_scf184961.g16 PE=4 SV=1 - [A0A060SAS1_PYCCI] | 27,14 | 4,39 | 1 | 3 | 3 | 8 | 1093 | 117,8 | 6,05 |
| A0A060SPR5 | Uncharacterized protein OS=Pycnoporus cinnabarinus GN=BN946_scf184880.g2 PE=4 SV=1 - [A0A060SPR5_PYCCI] | 25,61 | 13,53 | 1 | 6 | 6 | 8 | 658 | 72,4 | 5,10 |
| A0A060SDX7 | Glycoside Hydrolase Family 3 protein OS=Pycnoporus cinnabarinus GN=BN946_scf184977.g100 PE=4 SV=1 - [A0A060SDX7_PYCCI] | 24,26 | 9,30 | 1 | 4 | 4 | 8 | 656 | 69,6 | 4,97 |
| D2D3A8 | Laccase (Fragment) OS=Trametes punicea PE=4 SV=1 - [D2D3A8_9APHY] | 22,05 | 16,75 | 5 | 1 | 3 | 7 | 394 | 43,1 | 5,49 |
| A0A060SK35 | Glycoside Hydrolase Family 31 protein OS=Pycnoporus cinnabarinus GN=BN946_scf185022.g7 PE=3 SV=1 - [A0A060SK35_PYCCI] | 21,94 | 8,17 | 1 | 4 | 4 | 7 | 942 | 103,9 | 5,64 |
| Q7Z8Q1 | Lacasse (Fragment) OS=Trametes sanguinea PE=4 SV=1 - [Q7Z8Q1_9APHY] | 21,47 | 30,21 | 2 | 1 | 4 | 7 | 288 | 31,1 | 6,46 |

| | | | | | | | | | | |
|---|---|---|---|---|---|---|---|---|---|---|
| A0A060SSH9 | Uncharacterized protein OS=Pycnoporus cinnabarinus GN=BN946_scf184866.g20 PE=4 SV=1 - [A0A060SSH9_PYCCI] | 20,82 | 13,17 | 2 | 3 | 3 | 7 | 334 | 35,5 | 4,60 |
| A0A060SXH6 | Glyoxal oxidase OS=Pycnoporus cinnabarinus GN=BN946_scf184747.g41 PE=4 SV=1 - [A0A060SXH6_PYCCI] | 19,79 | 9,50 | 1 | 1 | 4 | 8 | 558 | 59,6 | 5,50 |
| A0A060S4R6 | Uncharacterized protein OS=Pycnoporus cinnabarinus GN=BN946_scf184817.g2 PE=4 SV=1 - [A0A060S4R6_PYCCI] | 18,54 | 16,95 | 1 | 4 | 4 | 7 | 354 | 38,7 | 5,47 |
| A0A060S3R1 | Alpha-amylase OS=Pycnoporus cinnabarinus GN=BN946_scf184992.g11 PE=3 SV=1 - [A0A060S3R1_PYCCI] | 18,38 | 5,05 | 1 | 3 | 3 | 6 | 634 | 67,4 | 4,88 |
| A0A060SRQ0 | Peptide hydrolase OS=Pycnoporus cinnabarinus GN=BN946_scf184473.g9 PE=3 SV=1 - [A0A060SRQ0_PYCCI] | 18,18 | 11,13 | 1 | 3 | 3 | 6 | 485 | 52,0 | 5,33 |
| A0A060S8Z6 | Glycoside Hydrolase Family 115 protein OS=Pycnoporus cinnabarinus GN=BN946_scf184989.g20 PE=4 SV=1 - [A0A060S8Z6_PYCCI] | 18,10 | 8,18 | 1 | 6 | 6 | 7 | 648 | 73,1 | 5,88 |
| A0A060SCQ2 | Copper radical oxidase OS=Pycnoporus cinnabarinus GN=BN946_scf185009.g14 PE=4 SV=1 - [A0A060SCQ2_PYCCI] | 17,81 | 3,82 | 1 | 2 | 2 | 5 | 996 | 106,4 | 4,48 |
| A0A060SIX2 | Beta-xylanase OS=Pycnoporus cinnabarinus GN=BN946_scf184449.g5 PE=3 SV=1 - [A0A060SIX2_PYCCI] | 17,39 | 15,28 | 1 | 3 | 3 | 6 | 301 | 32,1 | 4,92 |
| A0A060S827 | Peroxidase OS=Pycnoporus cinnabarinus GN=BN946_scf184569.g58 PE=3 SV=1 - [A0A060S827_PYCCI] | 16,85 | 7,97 | 1 | 1 | 4 | 7 | 364 | 38,7 | 4,83 |
| A0A060S784 | Uncharacterized protein OS=Pycnoporus cinnabarinus GN=BN946_scf184806.g23 PE=4 SV=1 - [A0A060S784_PYCCI] | 16,33 | 13,86 | 1 | 5 | 5 | 7 | 368 | 39,4 | 5,44 |
| A0A060SSA3 | Carbohydrate-Binding Module Family 1 / Glycoside Hydrolase Family 131 protein OS=Pycnoporus cinnabarinus GN=BN946_scf184654.g6 PE=4 SV=1 - [A0A060SSA3_PYCCI] | 15,96 | 9,61 | 1 | 2 | 2 | 5 | 281 | 30,2 | 4,82 |
| A0A060S8N9 | Uncharacterized protein OS=Pycnoporus cinnabarinus GN=BN946_scf184801.g14 PE=4 SV=1 - [A0A060S8N9_PYCCI] | 15,81 | 2,91 | 1 | 1 | 1 | 5 | 515 | 54,2 | 4,56 |
| A0A060SUU9 | Carbohydrate-Binding Module Family 1 / Glycoside Hydrolase Family 5 protein OS=Pycnoporus cinnabarinus GN=BN946_scf184727.g5 PE=3 SV=1 - [A0A060SUU9_PYCCI] | 15,02 | 9,33 | 2 | 2 | 2 | 5 | 386 | 40,6 | 5,05 |
| A0A060SLR9 | Iron reductase domain / Cellobiose dehydrogenase OS=Pycnoporus cinnabarinus GN=BN946_scf185013.g1 PE=3 SV=1 - [A0A060SLR9_PYCCI] | 14,45 | 6,76 | 2 | 1 | 5 | 7 | 1361 | 144,8 | 6,30 |

| | | | | | | | | | | |
|---|---|---|---|---|---|---|---|---|---|---|
| A0A060S8Q0 | Uncharacterized protein OS=Pycnoporus cinnabarinus GN=BN946_scf184748.g29 PE=4 SV=1 - [A0A060S8Q0_PYCCI] | 14,26 | 10,29 | 1 | 3 | 3 | 5 | 379 | 41,2 | 6,60 |
| A0A060S7X8 | Uncharacterized protein OS=Pycnoporus cinnabarinus GN=BN946_scf184843.g5 PE=4 SV=1 - [A0A060S7X8_PYCCI] | 14,07 | 14,81 | 1 | 2 | 2 | 4 | 135 | 15,2 | 5,74 |
| Q2TL94 | Tyrosinase OS=Trametes sanguinea GN=Tyr PE=2 SV=1 - [Q2TL94_9APHY] | 13,87 | 6,96 | 1 | 3 | 3 | 4 | 618 | 68,0 | 5,85 |
| A0A060SPP7 | Uncharacterized protein OS=Pycnoporus cinnabarinus GN=BN946_scf184867.g6 PE=4 SV=1 - [A0A060SPP7_PYCCI] | 13,68 | 9,48 | 1 | 3 | 3 | 5 | 443 | 47,3 | 6,23 |
| A0A060T0J0 | Carbohydrate-Binding Module Family 1 protein OS=Pycnoporus cinnabarinus GN=BN946_scf184616.g3 PE=4 SV=1 - [A0A060T0J0_PYCCI] | 13,41 | 2,84 | 1 | 2 | 2 | 5 | 703 | 76,7 | 8,28 |
| A0A060S548 | Glycoside Hydrolase Family 79 protein OS=Pycnoporus cinnabarinus GN=BN946_scf184817.g30 PE=4 SV=1 - [A0A060S548_PYCCI] | 13,23 | 6,06 | 1 | 2 | 2 | 5 | 396 | 42,0 | 7,24 |
| A0A060S956 | Glycoside Hydrolase Family 5 protein OS=Pycnoporus cinnabarinus GN=BN946_scf185000.g66 PE=3 SV=1 - [A0A060S956_PYCCI] | 13,22 | 37,62 | 1 | 3 | 3 | 3 | 210 | 23,1 | 6,24 |
| A0A060S8V2 | Uncharacterized protein OS=Pycnoporus cinnabarinus GN=BN946_scf184798.g90 PE=3 SV=1 - [A0A060S8V2_PYCCI] | 13,15 | 9,96 | 1 | 4 | 4 | 5 | 452 | 48,3 | 6,87 |
| A0A060S5D1 | Uncharacterized protein OS=Pycnoporus cinnabarinus GN=BN946_scf184961.g13 PE=3 SV=1 - [A0A060S5D1_PYCCI] | 12,99 | 5,44 | 1 | 2 | 2 | 4 | 478 | 52,6 | 4,97 |
| A0A060S5J2 | Glycoside Hydrolase Family 92 protein OS=Pycnoporus cinnabarinus GN=BN946_scf184817.g11 PE=4 SV=1 - [A0A060S5J2_PYCCI] | 12,97 | 8,63 | 1 | 4 | 4 | 5 | 707 | 78,2 | 5,40 |
| A0A060SBF2 | Uncharacterized protein OS=Pycnoporus cinnabarinus GN=BN946_scf184915.g29 PE=4 SV=1 - [A0A060SBF2_PYCCI] | 12,75 | 3,78 | 1 | 1 | 1 | 4 | 635 | 66,8 | 4,93 |
| A0A060SAZ4 | Glycoside Hydrolase Family 3 protein OS=Pycnoporus cinnabarinus GN=BN946_scf184817.g14 PE=4 SV=1 - [A0A060SAZ4_PYCCI] | 12,03 | 2,24 | 1 | 2 | 2 | 5 | 759 | 82,0 | 5,00 |
| A0A060S9U7 | Glycoside Hydrolase Family 18 / Carbohydrate-Binding Module Family 5 protein OS=Pycnoporus cinnabarinus GN=BN946_scf184992.g8 PE=3 SV=1 - [A0A060S9U7_PYCCI] | 11,32 | 4,68 | 1 | 3 | 3 | 4 | 534 | 57,6 | 5,14 |
| A0A060S3D6 | Glycoside Hydrolase Family 5 protein OS=Pycnoporus cinnabarinus GN=BN946_scf185000.g65 PE=4 SV=1 - [A0A060S3D6_PYCCI] | 11,26 | 18,50 | 1 | 3 | 3 | 4 | 200 | 22,3 | 5,83 |
| A0A060SSP9 | Glycoside Hydrolase Family 76 protein OS=Pycnoporus cinnabarinus GN=BN946_scf184926.g4 PE=4 SV=1 - [A0A060SSP9_PYCCI] | 11,25 | 6,94 | 1 | 1 | 1 | 3 | 173 | 18,9 | 4,91 |

| | | | | | | | | | | |
|---|---|---|---|---|---|---|---|---|---|---|
| A0A060S1Q5 | Pyranose oxidase OS=Pycnoporus cinnabarinus GN=BN946_scf184913.g15 PE=4 SV=1 - [A0A060S1Q5_PYCCI] | 11,20 | 9,65 | 1 | 5 | 5 | 5 | 715 | 79,2 | 8,75 |
| A0A060SZQ9 | Carboxypeptidase OS=Pycnoporus cinnabarinus GN=BN946_scf184826.g3 PE=3 SV=1 - [A0A060SZQ9_PYCCI] | 10,51 | 7,16 | 1 | 3 | 3 | 4 | 489 | 52,8 | 5,06 |
| E1A4C3 | Peroxidase (Fragment) OS=Pycnoporus cinnabarinus GN=mnp2 PE=3 SV=1 - [E1A4C3_PYCCI] | 10,12 | 12,44 | 1 | 1 | 4 | 5 | 201 | 21,4 | 4,51 |
| A0A060S604 | Glycoside Hydrolase Family 43 protein OS=Pycnoporus cinnabarinus GN=BN946_scf184884.g56 PE=4 SV=1 - [A0A060S604_PYCCI] | 10,09 | 9,04 | 1 | 1 | 1 | 3 | 177 | 18,9 | 4,46 |
| A0A060SYU5 | Glycoside Hydrolase Family 92 protein OS=Pycnoporus cinnabarinus GN=BN946_scf184969.g74 PE=4 SV=1 - [A0A060SYU5_PYCCI] | 10,01 | 5,60 | 1 | 3 | 3 | 4 | 661 | 73,5 | 5,40 |
| A0A060SH23 | Auxilliary Activities Family 9 / Carbohydrate-Binding Module Family 1 protein OS=Pycnoporus cinnabarinus GN=BN946_scf184915.g57 PE=4 SV=1 - [A0A060SH23_PYCCI] | 9,64 | 4,97 | 1 | 1 | 1 | 3 | 322 | 33,7 | 5,31 |
| A0A060SKG6 | Uncharacterized protein OS=Pycnoporus cinnabarinus GN=BN946_scf185002.g93 PE=4 SV=1 - [A0A060SKG6_PYCCI] | 9,52 | 4,66 | 1 | 4 | 4 | 4 | 580 | 62,5 | 5,14 |
| A0A060SJ44 | Uncharacterized protein OS=Pycnoporus cinnabarinus GN=BN946_scf185043.g258 PE=4 SV=1 - [A0A060SJ44_PYCCI] | 9,45 | 18,88 | 1 | 2 | 2 | 3 | 143 | 14,8 | 5,68 |
| A0A060SP82 | Carbohydrate Esterase Family 16 protein OS=Pycnoporus cinnabarinus GN=BN946_scf184751.g9 PE=4 SV=1 - [A0A060SP82_PYCCI] | 9,07 | 6,38 | 1 | 1 | 1 | 3 | 345 | 37,0 | 4,75 |
| A0A060S9M8 | Uncharacterized protein OS=Pycnoporus cinnabarinus GN=BN946_scf184777.g14 PE=4 SV=1 - [A0A060S9M8_PYCCI] | 8,76 | 2,52 | 1 | 3 | 3 | 4 | 913 | 100,1 | 5,87 |
| A0A060S792 | Glycoside Hydrolase Family 12 protein OS=Pycnoporus cinnabarinus GN=BN946_scf184806.g33 PE=3 SV=1 - [A0A060S792_PYCCI] | 7,76 | 9,69 | 1 | 2 | 2 | 3 | 258 | 27,0 | 5,08 |
| A0A060SLF1 | Glycoside Hydrolase Family 3 protein OS=Pycnoporus cinnabarinus GN=BN946_scf185007.g108 PE=4 SV=1 - [A0A060SLF1_PYCCI] | 7,73 | 4,10 | 1 | 3 | 3 | 3 | 757 | 81,6 | 5,52 |
| A0A060SCG5 | Glycoside Hydrolase Family 20 protein OS=Pycnoporus cinnabarinus GN=BN946_scf184970.g39 PE=4 SV=1 - [A0A060SCG5_PYCCI] | 7,57 | 5,30 | 1 | 2 | 2 | 3 | 415 | 45,6 | 5,43 |
| A0A060S7B6 | Carbohydrate Esterase Family 15 protein OS=Pycnoporus cinnabarinus GN=BN946_scf184856.g1 PE=4 SV=1 - [A0A060S7B6_PYCCI] | 7,51 | 6,94 | 1 | 2 | 2 | 3 | 418 | 45,8 | 5,69 |
| A0A060SAA4 | Glycoside Hydrolase Family 18 protein OS=Pycnoporus cinnabarinus GN=BN946_scf184845.g46 PE=3 SV=1 - [A0A060SAA4_PYCCI] | 7,40 | 5,08 | 1 | 2 | 2 | 3 | 453 | 47,0 | 4,35 |

| | | | | | | | | | | |
|---|---|---|---|---|---|---|---|---|---|---|
| A0A060SQW9 | Auxilliary Activities Family 9 protein OS=Pycnoporus cinnabarinus GN=BN946_scf184831.g7 PE=4 SV=1 - [A0A060SQW9_PYCCI] | 7,29 | 7,51 | 1 | 1 | 1 | 2 | 253 | 26,7 | 7,02 |
| A0A060SZR2 | Uncharacterized protein OS=Pycnoporus cinnabarinus GN=BN946_scf184993.g7 PE=4 SV=1 - [A0A060SZR2_PYCCI] | 7,29 | 11,53 | 1 | 2 | 2 | 2 | 347 | 37,5 | 5,38 |
| A0A060SVL6 | Uncharacterized protein OS=Pycnoporus cinnabarinus GN=BN946_scf184750.g8 PE=3 SV=1 - [A0A060SVL6_PYCCI] | 7,08 | 5,42 | 1 | 1 | 1 | 2 | 757 | 82,3 | 7,09 |
| A0A060SLC0 | Uncharacterized protein OS=Pycnoporus cinnabarinus GN=BN946_scf184945.g48 PE=4 SV=1 - [A0A060SLC0_PYCCI] | 6,91 | 2,73 | 1 | 1 | 1 | 3 | 513 | 52,1 | 5,43 |
| A0A060SHS2 | Glycoside Hydrolase Family 125 protein OS=Pycnoporus cinnabarinus GN=BN946_scf185043.g112 PE=4 SV=1 - [A0A060SHS2_PYCCI] | 6,91 | 4,48 | 1 | 1 | 1 | 2 | 469 | 51,8 | 6,18 |
| A0A060SBL7 | Carboxypeptidase OS=Pycnoporus cinnabarinus GN=BN946_scf184766.g13 PE=3 SV=1 - [A0A060SBL7_PYCCI] | 6,69 | 3,22 | 1 | 2 | 2 | 3 | 653 | 71,3 | 5,53 |
| A0A060S2S5 | Glycoside Hydrolase Family 28 protein OS=Pycnoporus cinnabarinus GN=BN946_scf184996.g46 PE=3 SV=1 - [A0A060S2S5_PYCCI] | 6,66 | 4,22 | 1 | 1 | 1 | 2 | 379 | 39,3 | 5,31 |
| A0A060SXP7 | Uncharacterized protein OS=Pycnoporus cinnabarinus GN=BN946_scf184380.g10 PE=4 SV=1 - [A0A060SXP7_PYCCI] | 6,66 | 4,74 | 1 | 2 | 2 | 3 | 527 | 58,1 | 5,30 |
| A0A060SB47 | Auxilliary Activities Family 9 protein OS=Pycnoporus cinnabarinus GN=BN946_scf184915.g56 PE=4 SV=1 - [A0A060SB47_PYCCI] | 6,56 | 1,74 | 1 | 2 | 2 | 3 | 922 | 101,0 | 7,65 |
| A0A060STA0 | Uncharacterized protein OS=Pycnoporus cinnabarinus GN=BN946_scf184993.g25 PE=4 SV=1 - [A0A060STA0_PYCCI] | 6,35 | 4,81 | 1 | 1 | 1 | 2 | 291 | 31,4 | 5,80 |
| A0A060SXD3 | Glycoside Hydrolase Family 18 protein OS=Pycnoporus cinnabarinus GN=BN946_scf184747.g1 PE=3 SV=1 - [A0A060SXD3_PYCCI] | 6,30 | 2,64 | 1 | 2 | 2 | 2 | 568 | 60,9 | 6,47 |
| A0A060SSJ5 | Carbohydrate Esterase Family 16 protein OS=Pycnoporus cinnabarinus GN=BN946_scf184916.g7 PE=4 SV=1 - [A0A060SSJ5_PYCCI] | 6,20 | 3,91 | 1 | 1 | 1 | 2 | 358 | 40,1 | 5,92 |
| A0A060SQK1 | Glycoside Hydrolase Family 95 protein OS=Pycnoporus cinnabarinus GN=BN946_scf184593.g5 PE=4 SV=1 - [A0A060SQK1_PYCCI] | 6,10 | 1,62 | 1 | 1 | 1 | 2 | 677 | 73,8 | 5,02 |
| A0A060SEI1 | Glycoside Hydrolase Family 55 protein OS=Pycnoporus cinnabarinus GN=BN946_scf185002.g87 PE=4 SV=1 - [A0A060SEI1_PYCCI] | 6,05 | 3,52 | 1 | 2 | 2 | 2 | 768 | 80,6 | 5,02 |
| A0A060S4G5 | Copper radical oxidase OS=Pycnoporus cinnabarinus GN=BN946_scf184992.g45 PE=4 SV=1 - [A0A060S4G5_PYCCI] | 5,52 | 2,16 | 1 | 1 | 1 | 2 | 740 | 78,0 | 5,30 |

| | | | | | | | | | | |
|---|---|---|---|---|---|---|---|---|---|---|
| A0A060SYQ9 | Uncharacterized protein OS=Pycnoporus cinnabarinus GN=BN946_scf184787.g18 PE=3 SV=1 - [A0A060SYQ9_PYCCI] | 5,38 | 2,65 | 1 | 1 | 1 | 2 | 415 | 43,1 | 4,44 |
| A0A060S4J3 | Uncharacterized protein OS=Pycnoporus cinnabarinus GN=BN946_scf185042.g152 PE=4 SV=1 - [A0A060S4J3_PYCCI] | 5,30 | 6,92 | 1 | 1 | 1 | 2 | 318 | 33,9 | 5,41 |
| A0A060SAF4 | Uncharacterized protein OS=Pycnoporus cinnabarinus GN=BN946_scf184908.g52 PE=4 SV=1 - [A0A060SAF4_PYCCI] | 5,28 | 1,21 | 1 | 1 | 1 | 2 | 661 | 71,9 | 5,69 |
| A0A060S724 | Carbohydrate Esterase Family 16 protein OS=Pycnoporus cinnabarinus GN=BN946_scf184938.g40 PE=4 SV=1 - [A0A060S724_PYCCI] | 5,27 | 3,45 | 1 | 1 | 1 | 2 | 319 | 34,7 | 5,54 |
| A0A060S3L6 | Uncharacterized protein OS=Pycnoporus cinnabarinus GN=BN946_scf185000.g33 PE=4 SV=1 - [A0A060S3L6_PYCCI] | 5,19 | 5,37 | 1 | 2 | 2 | 2 | 410 | 44,2 | 7,43 |
| A0A060SCR0 | Glycoside Hydrolase Family 12 protein OS=Pycnoporus cinnabarinus GN=BN946_scf184970.g119 PE=3 SV=1 - [A0A060SCR0_PYCCI] | 5,18 | 4,20 | 1 | 1 | 1 | 2 | 262 | 28,1 | 4,96 |
| A0A060SQD4 | Carboxypeptidase OS=Pycnoporus cinnabarinus GN=BN946_scf185011.g20 PE=3 SV=1 - [A0A060SQD4_PYCCI] | 4,68 | 6,58 | 1 | 1 | 2 | 2 | 395 | 43,8 | 5,30 |
| A0A060SE18 | Glycoside Hydrolase Family 78 protein OS=Pycnoporus cinnabarinus GN=BN946_scf184977.g152 PE=4 SV=1 - [A0A060SE18_PYCCI] | 4,48 | 3,88 | 1 | 1 | 1 | 1 | 670 | 72,1 | 5,57 |
| A0A060SE28 | Uncharacterized protein OS=Pycnoporus cinnabarinus GN=BN946_scf184985.g44 PE=4 SV=1 - [A0A060SE28_PYCCI] | 4,32 | 5,88 | 2 | 1 | 1 | 2 | 136 | 15,3 | 6,95 |
| A0A060SP89 | Glycoside Hydrolase Family 28 protein OS=Pycnoporus cinnabarinus GN=BN946_scf184414.g9 PE=3 SV=1 - [A0A060SP89_PYCCI] | 4,26 | 4,38 | 1 | 2 | 2 | 2 | 434 | 45,8 | 6,39 |
| A0A060SY86 | Auxilliary Activities Family 9 / Carbohydrate-Binding Module Family 1 protein OS=Pycnoporus cinnabarinus GN=BN946_scf184747.g17 PE=4 SV=1 - [A0A060SY86_PYCCI] | 4,10 | 6,31 | 1 | 1 | 1 | 1 | 317 | 33,0 | 4,74 |
| A0A060SK99 | Uncharacterized protein (Fragment) OS=Pycnoporus cinnabarinus GN=BN946_scf184985.g88 PE=4 SV=1 - [A0A060SK99_PYCCI] | 3,89 | 1,73 | 1 | 1 | 1 | 1 | 752 | 80,9 | 5,68 |
| A0A060SNI3 | Uncharacterized protein OS=Pycnoporus cinnabarinus GN=BN946_scf184649.g23 PE=4 SV=1 - [A0A060SNI3_PYCCI] | 3,60 | 3,64 | 1 | 1 | 1 | 2 | 990 | 109,8 | 6,92 |
| A0A060SMQ5 | Uncharacterized protein OS=Pycnoporus cinnabarinus GN=BN946_scf185013.g154 PE=3 SV=1 - [A0A060SMQ5_PYCCI] | 3,58 | 2,00 | 1 | 1 | 1 | 1 | 550 | 59,7 | 5,29 |
| A0A060S8R4 | Uncharacterized protein OS=Pycnoporus cinnabarinus GN=BN946_scf184652.g1 PE=4 SV=1 - [A0A060S8R4_PYCCI] | 3,57 | 13,95 | 1 | 1 | 1 | 1 | 294 | 32,5 | 8,65 |

| | | | | | | | | | | |
|---|---|---|---|---|---|---|---|---|---|---|
| A0A060S811 | Uncharacterized protein OS=Pycnoporus cinnabarinus GN=BN946_scf184613.g6 PE=3 SV=1 - [A0A060S811_PYCCI] | 3,45 | 7,77 | 1 | 1 | 1 | 2 | 373 | 41,5 | 5,68 |
| Q5TLD0 | Glyceraldehyde-3-phosphate dehydrogenase OS=Pycnoporus coccineus GN=gpd PE=3 SV=1 - [Q5TLD0_PYCCO] | 3,44 | 4,13 | 2 | 1 | 1 | 1 | 339 | 36,3 | 7,44 |
| A0A060SEP2 | Uncharacterized protein OS=Pycnoporus cinnabarinus GN=BN946_scf184994.g53 PE=4 SV=1 - [A0A060SEP2_PYCCI] | 3,33 | 4,21 | 1 | 1 | 1 | 1 | 285 | 30,5 | 4,63 |
| A0A060SHI1 | Uncharacterized protein OS=Pycnoporus cinnabarinus GN=BN946_scf184915.g38 PE=4 SV=1 - [A0A060SHI1_PYCCI] | 3,32 | 8,51 | 1 | 1 | 1 | 2 | 423 | 46,2 | 8,68 |
| A0A060SWE9 | Beta-hexosaminidase OS=Pycnoporus cinnabarinus GN=BN946_scf184978.g17 PE=3 SV=1 - [A0A060SWE9_PYCCI] | 3,30 | 6,67 | 1 | 1 | 1 | 1 | 570 | 61,7 | 5,88 |
| A0A060S704 | Uncharacterized protein OS=Pycnoporus cinnabarinus GN=BN946_scf184942.g47 PE=3 SV=1 - [A0A060S704_PYCCI] | 3,30 | 2,37 | 1 | 1 | 1 | 1 | 548 | 58,8 | 5,08 |
| A0A060SPL9 | Uncharacterized protein OS=Pycnoporus cinnabarinus GN=BN946_scf184622.g5 PE=4 SV=1 - [A0A060SPL9_PYCCI] | 3,22 | 5,88 | 1 | 1 | 1 | 1 | 255 | 27,6 | 4,83 |
| A0A060SUL2 | Glycoside Hydrolase Family 18 protein OS=Pycnoporus cinnabarinus GN=BN946_scf184688.g4 PE=3 SV=1 - [A0A060SUL2_PYCCI] | 3,21 | 3,59 | 1 | 1 | 1 | 1 | 334 | 35,2 | 4,75 |
| A0A060SB04 | Polysaccharide Lyase Family 8 protein OS=Pycnoporus cinnabarinus GN=BN946_scf184911.g5 PE=4 SV=1 - [A0A060SB04_PYCCI] | 3,20 | 1,44 | 1 | 1 | 1 | 1 | 764 | 81,4 | 5,03 |
| A0A060SI63 | Uncharacterized protein OS=Pycnoporus cinnabarinus GN=BN946_scf184962.g71 PE=4 SV=1 - [A0A060SI63_PYCCI] | 3,20 | 4,98 | 1 | 1 | 1 | 1 | 261 | 27,5 | 4,87 |
| A0A060SB69 | Uncharacterized protein OS=Pycnoporus cinnabarinus GN=BN946_scf184911.g70 PE=3 SV=1 - [A0A060SB69_PYCCI] | 3,18 | 5,45 | 1 | 1 | 1 | 1 | 422 | 43,1 | 6,47 |
| A0A060T147 | Uncharacterized protein OS=Pycnoporus cinnabarinus GN=BN946_scf184991.g2 PE=4 SV=1 - [A0A060T147_PYCCI] | 3,11 | 4,44 | 1 | 1 | 1 | 1 | 901 | 100,8 | 7,49 |
| A0A060STT9 | Glycoside Hydrolase Family 76 protein OS=Pycnoporus cinnabarinus GN=BN946_scf184926.g3 PE=4 SV=1 - [A0A060STT9_PYCCI] | 3,09 | 4,52 | 1 | 1 | 1 | 1 | 221 | 24,2 | 4,97 |
| A0A060SNY9 | Uncharacterized protein OS=Pycnoporus cinnabarinus GN=BN946_scf184833.g2 PE=4 SV=1 - [A0A060SNY9_PYCCI] | 3,08 | 1,66 | 1 | 1 | 1 | 1 | 964 | 105,2 | 5,60 |
| A0A060S9S1 | Uncharacterized protein OS=Pycnoporus cinnabarinus GN=BN946_scf184830.g2 PE=4 SV=1 - [A0A060S9S1_PYCCI] | 3,00 | 1,74 | 1 | 1 | 1 | 1 | 2127 | 234,3 | 7,64 |

| | | | | | | | | | | |
|---|---|---|---|---|---|---|---|---|---|---|
| A0A060S8V8 | Uncharacterized protein OS=Pycnoporus cinnabarinus GN=BN946_scf184829.g40 PE=4 SV=1 - [A0A060S8V8_PYCCI] | 2,99 | 4,08 | 1 | 1 | 1 | 1 | 981 | 105,9 | 7,27 |
| A0A060S4J1 | Aryl-alcohol oxidase OS=Pycnoporus cinnabarinus GN=BN946_scf184746.g13 PE=3 SV=1 - [A0A060S4J1_PYCCI] | 2,98 | 3,04 | 1 | 1 | 1 | 1 | 460 | 49,3 | 5,94 |
| A0A060SJV2 | Carbohydrate-Binding Module Family 1 / Carbohydrate Esterase Family 1 protein OS=Pycnoporus cinnabarinus GN=BN946_scf184980.g45 PE=4 SV=1 - [A0A060SJV2_PYCCI] | 2,96 | 3,36 | 1 | 1 | 1 | 1 | 357 | 37,6 | 5,92 |
| A0A060SPD3 | Glycoside Hydrolase Family 18 protein OS=Pycnoporus cinnabarinus GN=BN946_scf184665.g23 PE=3 SV=1 - [A0A060SPD3_PYCCI] | 2,93 | 2,05 | 1 | 1 | 1 | 1 | 391 | 41,8 | 4,93 |
| A0A060SPM8 | Uncharacterized protein OS=Pycnoporus cinnabarinus GN=BN946_scf184361.g6 PE=4 SV=1 - [A0A060SPM8_PYCCI] | 2,93 | 3,49 | 1 | 1 | 1 | 2 | 1260 | 137,1 | 9,39 |
| A0A060SYY4 | Uncharacterized protein OS=Pycnoporus cinnabarinus GN=BN946_scf184993.g21 PE=4 SV=1 - [A0A060SYY4_PYCCI] | 2,89 | 11,69 | 1 | 1 | 1 | 1 | 385 | 44,6 | 6,81 |
| A0A060S881 | Uncharacterized protein OS=Pycnoporus cinnabarinus GN=BN946_scf184996.g24 PE=4 SV=1 - [A0A060S881_PYCCI] | 2,89 | 5,19 | 1 | 1 | 1 | 1 | 501 | 55,7 | 9,47 |
| A0A060SDN1 | Uncharacterized protein OS=Pycnoporus cinnabarinus GN=BN946_scf184999.g9 PE=4 SV=1 - [A0A060SDN1_PYCCI] | 2,86 | 8,18 | 1 | 1 | 1 | 1 | 110 | 12,2 | 9,25 |
| A0A060SQG8 | Uncharacterized protein OS=Pycnoporus cinnabarinus GN=BN946_scf184868.g39 PE=4 SV=1 - [A0A060SQG8_PYCCI] | 2,85 | 3,30 | 1 | 1 | 1 | 1 | 696 | 76,2 | 6,58 |
| A0A060SES6 | Uncharacterized protein OS=Pycnoporus cinnabarinus GN=BN946_scf185007.g76 PE=4 SV=1 - [A0A060SES6_PYCCI] | 2,82 | 1,32 | 1 | 1 | 1 | 1 | 1663 | 184,8 | 5,44 |
| A0A060SF67 | Uncharacterized protein OS=Pycnoporus cinnabarinus GN=BN946_scf184804.g11 PE=4 SV=1 - [A0A060SF67_PYCCI] | 2,79 | 4,51 | 1 | 1 | 1 | 1 | 288 | 30,6 | 5,00 |
| A0A060SM02 | Uncharacterized protein OS=Pycnoporus cinnabarinus GN=BN946_scf184883.g14 PE=4 SV=1 - [A0A060SM02_PYCCI] | 2,77 | 11,83 | 1 | 1 | 1 | 1 | 355 | 37,2 | 4,79 |
| A0A060SBS6 | Uncharacterized protein OS=Pycnoporus cinnabarinus GN=BN946_scf184943.g1 PE=4 SV=1 - [A0A060SBS6_PYCCI] | 2,74 | 15,70 | 1 | 1 | 1 | 1 | 242 | 27,7 | 6,65 |
| A0A060SDI3 | Uncharacterized protein OS=Pycnoporus cinnabarinus GN=BN946_scf184983.g60 PE=4 SV=1 - [A0A060SDI3_PYCCI] | 2,73 | 3,30 | 1 | 1 | 1 | 1 | 303 | 32,5 | 5,49 |
| A0A060SR95 | Uncharacterized protein OS=Pycnoporus cinnabarinus GN=BN946_scf184901.g12 PE=4 SV=1 - [A0A060SR95_PYCCI] | 2,73 | 3,55 | 1 | 1 | 1 | 1 | 648 | 71,9 | 5,34 |

| | | | | | | | | | | |
|---|---|---|---|---|---|---|---|---|---|---|
| A0A060SMW0 | Uncharacterized protein OS=Pycnoporus cinnabarinus GN=BN946_scf184951.g20 PE=4 SV=1 - [A0A060SMW0_PYCCI] | 2,69 | 2,68 | 1 | 1 | 1 | 1 | 672 | 74,7 | 9,14 |
| A0A060SAD3 | Uncharacterized protein OS=Pycnoporus cinnabarinus GN=BN946_scf185042.g151 PE=4 SV=1 - [A0A060SAD3_PYCCI] | 2,68 | 2,81 | 1 | 1 | 1 | 1 | 570 | 59,1 | 4,72 |
| A0A060STF1 | Uncharacterized protein OS=Pycnoporus cinnabarinus GN=BN946_scf184857.g12 PE=4 SV=1 - [A0A060STF1_PYCCI] | 2,68 | 2,75 | 1 | 1 | 1 | 1 | 436 | 46,9 | 5,17 |
| A0A060SIS0 | Glycoside Hydrolase Family 16 protein OS=Pycnoporus cinnabarinus GN=BN946_scf184392.g3 PE=4 SV=1 - [A0A060SIS0_PYCCI] | 2,62 | 1,56 | 1 | 1 | 1 | 1 | 707 | 77,5 | 4,91 |
| A0A060SMK5 | Uncharacterized protein OS=Pycnoporus cinnabarinus GN=BN946_scf185013.g94 PE=4 SV=1 - [A0A060SMK5_PYCCI] | 2,58 | 8,87 | 1 | 1 | 1 | 1 | 124 | 13,7 | 4,87 |
| A0A060SN13 | Uncharacterized protein OS=Pycnoporus cinnabarinus GN=BN946_scf184858.g26 PE=3 SV=1 - [A0A060SN13_PYCCI] | 2,56 | 2,43 | 1 | 1 | 1 | 1 | 412 | 44,9 | 4,78 |
| A0A060SCZ8 | Uncharacterized protein OS=Pycnoporus cinnabarinus GN=BN946_scf184942.g53 PE=4 SV=1 - [A0A060SCZ8_PYCCI] | 2,55 | 3,63 | 1 | 1 | 1 | 1 | 248 | 26,0 | 8,57 |
| A0A060SRX7 | Glycoside Hydrolase Family 30 protein OS=Pycnoporus cinnabarinus GN=BN946_scf184657.g15 PE=3 SV=1 - [A0A060SRX7_PYCCI] | 2,54 | 2,33 | 2 | 1 | 1 | 1 | 557 | 62,0 | 5,91 |
| Q4TWI5 | Elongation factor 1-alpha (Fragment) OS=Pycnoporus sp. ZW02.30 GN=tef1 PE=3 SV=1 - [Q4TWI5_9APHY] | 2,49 | 2,73 | 2 | 1 | 1 | 1 | 403 | 44,0 | 8,16 |
| A0A060SEI3 | Uncharacterized protein OS=Pycnoporus cinnabarinus GN=BN946_scf185002.g92 PE=4 SV=1 - [A0A060SEI3_PYCCI] | 2,48 | 1,40 | 1 | 1 | 1 | 1 | 643 | 70,7 | 7,42 |
| A0A060SFK1 | Uncharacterized protein OS=Pycnoporus cinnabarinus GN=BN946_scf185007.g40 PE=4 SV=1 - [A0A060SFK1_PYCCI] | 2,47 | 1,42 | 1 | 1 | 1 | 1 | 772 | 84,9 | 5,19 |
| A0A060SFJ4 | Uncharacterized protein OS=Pycnoporus cinnabarinus GN=BN946_scf184845.g38 PE=4 SV=1 - [A0A060SFJ4_PYCCI] | 2,42 | 7,24 | 1 | 1 | 1 | 1 | 304 | 34,7 | 5,41 |
| A0A060SQG0 | Uncharacterized protein OS=Pycnoporus cinnabarinus GN=BN946_scf184978.g1 PE=4 SV=1 - [A0A060SQG0_PYCCI] | 2,40 | 1,81 | 1 | 1 | 1 | 1 | 497 | 53,2 | 5,19 |
| A0A060S411 | Glycoside Hydrolase Family 18 / Carbohydrate-Binding Module Family 5 protein OS=Pycnoporus cinnabarinus GN=BN946_scf184992.g9 PE=4 SV=1 - [A0A060S411_PYCCI] | 2,36 | 2,21 | 1 | 1 | 1 | 1 | 634 | 67,5 | 8,90 |
| A0A060SR05 | Beta-glucosidase OS=Pycnoporus cinnabarinus GN=BN946_scf185004.g8 PE=3 SV=1 - [A0A060SR05_PYCCI] | 2,34 | 0,99 | 1 | 1 | 1 | 1 | 705 | 77,1 | 5,52 |

| | | | | | | | | | | |
|---|---|---|---|---|---|---|---|---|---|---|
| A0A060S4D9 | Uncharacterized protein OS=Pycnoporus cinnabarinus GN=BN946_scf184992.g10 PE=4 SV=1 - [A0A060S4D9_PYCCI] | 2,32 | 2,42 | 1 | 1 | 1 | 1 | 1034 | 116,8 | 6,76 |
| A0A060SHV5 | Uncharacterized protein OS=Pycnoporus cinnabarinus GN=BN946_scf185043.g147 PE=4 SV=1 - [A0A060SHV5_PYCCI] | 2,32 | 3,27 | 1 | 1 | 1 | 1 | 214 | 22,2 | 4,41 |
| A0A060SG20 | Uncharacterized protein OS=Pycnoporus cinnabarinus GN=BN946_scf185013.g102 PE=4 SV=1 - [A0A060SG20_PYCCI] | 2,32 | 3,63 | 1 | 1 | 1 | 1 | 908 | 98,1 | 9,96 |
| A0A060SI08 | Uncharacterized protein OS=Pycnoporus cinnabarinus GN=BN946_scf185043.g203 PE=4 SV=1 - [A0A060SI08_PYCCI] | 2,31 | 2,67 | 1 | 1 | 1 | 1 | 562 | 57,9 | 9,22 |
| A0A060SK22 | Uncharacterized protein OS=Pycnoporus cinnabarinus GN=BN946_scf184985.g3 PE=3 SV=1 - [A0A060SK22_PYCCI] | 2,27 | 2,31 | 1 | 1 | 1 | 1 | 390 | 41,7 | 4,91 |
| A0A060S9M1 | Uncharacterized protein OS=Pycnoporus cinnabarinus GN=BN946_scf184777.g4 PE=4 SV=1 - [A0A060S9M1_PYCCI] | 2,24 | 3,69 | 1 | 1 | 1 | 1 | 217 | 23,5 | 8,94 |
| A0A060SBW4 | Histone H3 OS=Pycnoporus cinnabarinus GN=BN946_scf184884.g22 PE=3 SV=1 - [A0A060SBW4_PYCCI] | 2,22 | 5,11 | 4 | 1 | 1 | 1 | 137 | 15,4 | 11,52 |
| A0A060SQY0 | Uncharacterized protein OS=Pycnoporus cinnabarinus GN=BN946_scf184950.g7 PE=4 SV=1 - [A0A060SQY0_PYCCI] | 2,21 | 3,77 | 1 | 1 | 1 | 1 | 212 | 23,5 | 6,21 |
| A0A060SC99 | Glycoside Hydrolase Family 18 protein OS=Pycnoporus cinnabarinus GN=BN946_scf184939.g35 PE=3 SV=1 - [A0A060SC99_PYCCI] | 2,20 | 2,63 | 1 | 1 | 1 | 1 | 419 | 45,3 | 5,06 |
| A0A060SIH6 | Uncharacterized protein OS=Pycnoporus cinnabarinus GN=BN946_scf185043.g257 PE=4 SV=1 - [A0A060SIH6_PYCCI] | 2,17 | 5,59 | 1 | 1 | 1 | 1 | 143 | 14,9 | 5,00 |
| A0A060SDL9 | Carboxylic ester hydrolase (Fragment) OS=Pycnoporus cinnabarinus GN=BN946_scf184569.g41 PE=3 SV=1 - [A0A060SDL9_PYCCI] | 2,16 | 1,35 | 1 | 1 | 1 | 1 | 592 | 63,4 | 5,77 |
| A0A060SB90 | Uncharacterized protein OS=Pycnoporus cinnabarinus GN=BN946_scf184851.g41 PE=4 SV=1 - [A0A060SB90_PYCCI] | 2,13 | 1,25 | 1 | 1 | 1 | 1 | 642 | 71,8 | 5,29 |
| A0A060S977 | Uncharacterized protein OS=Pycnoporus cinnabarinus GN=BN946_scf185000.g91 PE=4 SV=1 - [A0A060S977_PYCCI] | 2,12 | 2,51 | 1 | 1 | 1 | 1 | 438 | 47,3 | 9,50 |
| A0A060SME9 | Uncharacterized protein OS=Pycnoporus cinnabarinus GN=BN946_scf184585.g15 PE=4 SV=1 - [A0A060SME9_PYCCI] | 2,10 | 0,95 | 1 | 1 | 1 | 1 | 737 | 80,8 | 5,57 |
| A0A060SHF8 | Uncharacterized protein OS=Pycnoporus cinnabarinus GN=BN946_scf184939.g67 PE=4 SV=1 - [A0A060SHF8_PYCCI] | 2,09 | 0,77 | 1 | 1 | 1 | 1 | 1045 | 109,4 | 5,88 |

| | | | | | | | | | | |
|---|---|---|---|---|---|---|---|---|---|---|
| A0A060SNT0 | Uncharacterized protein OS=Pycnoporus cinnabarinus GN=BN946_scf184819.g2 PE=4 SV=1 - [A0A060SNT0_PYCCI] | 2,08 | 1,67 | 1 | 1 | 1 | 1 | 418 | 44,4 | 5,67 |
| A0A060ST21 | Peroxidase OS=Pycnoporus cinnabarinus GN=BN946_scf184969.g43 PE=3 SV=1 - [A0A060ST21_PYCCI] | 2,07 | 1,78 | 1 | 1 | 1 | 1 | 394 | 42,4 | 5,03 |
| A0A060S8N7 | Glycoside Hydrolase Family 115 protein OS=Pycnoporus cinnabarinus GN=BN946_scf184989.g9 PE=4 SV=1 - [A0A060S8N7_PYCCI] | 2,04 | 0,64 | 1 | 1 | 1 | 1 | 1087 | 120,2 | 5,19 |
| A0A060SFJ3 | Uncharacterized protein OS=Pycnoporus cinnabarinus GN=BN946_scf185007.g30 PE=4 SV=1 - [A0A060SFJ3_PYCCI] | 2,01 | 1,44 | 1 | 1 | 1 | 1 | 765 | 83,7 | 5,01 |
| A0A060SFE5 | Uncharacterized protein OS=Pycnoporus cinnabarinus GN=BN946_scf184844.g127 PE=4 SV=1 - [A0A060SFE5_PYCCI] | 2,00 | 2,03 | 1 | 1 | 1 | 1 | 394 | 39,2 | 4,79 |
| A0A060S5J9 | Auxilliary Activities Family 9 protein OS=Pycnoporus cinnabarinus GN=BN946_scf184817.g21 PE=4 SV=1 - [A0A060S5J9_PYCCI] | 1,99 | 2,04 | 1 | 1 | 1 | 1 | 737 | 81,0 | 7,58 |
| A0A060ST62 | Uncharacterized protein OS=Pycnoporus cinnabarinus GN=BN946_scf184855.g2 PE=4 SV=1 - [A0A060ST62_PYCCI] | 1,99 | 3,39 | 1 | 1 | 1 | 1 | 236 | 25,9 | 9,77 |
| A0A060S923 | Uncharacterized protein OS=Pycnoporus cinnabarinus GN=BN946_scf184844.g16 PE=4 SV=1 - [A0A060S923_PYCCI] | 1,96 | 11,89 | 1 | 1 | 1 | 1 | 143 | 15,4 | 4,91 |
| A0A060S404 | Glycoside Hydrolase Family 88 protein OS=Pycnoporus cinnabarinus GN=BN946_scf185000.g29 PE=4 SV=1 - [A0A060S404_PYCCI] | 1,96 | 6,84 | 1 | 1 | 1 | 1 | 190 | 21,1 | 8,60 |
| A0A060SX36 | Uncharacterized protein OS=Pycnoporus cinnabarinus GN=BN946_scf184978.g8 PE=4 SV=1 - [A0A060SX36_PYCCI] | 1,96 | 1,69 | 2 | 1 | 1 | 1 | 592 | 62,4 | 5,12 |
| A0A060SA81 | Glycoside Hydrolase Family 79 protein OS=Pycnoporus cinnabarinus GN=BN946_scf184908.g155 PE=4 SV=1 - [A0A060SA81_PYCCI] | 1,95 | 2,65 | 1 | 1 | 1 | 1 | 491 | 51,4 | 6,21 |
| A0A060SYD2 | Uncharacterized protein OS=Pycnoporus cinnabarinus GN=BN946_scf184926.g5 PE=4 SV=1 - [A0A060SYD2_PYCCI] | 1,94 | 5,26 | 1 | 1 | 1 | 1 | 418 | 45,3 | 8,38 |
| A0A060SBR5 | Carboxylic ester hydrolase OS=Pycnoporus cinnabarinus GN=BN946_scf184940.g99 PE=3 SV=1 - [A0A060SBR5_PYCCI] | 1,93 | 1,47 | 1 | 1 | 1 | 1 | 543 | 59,5 | 4,86 |
| A0A060SNK6 | Uncharacterized protein OS=Pycnoporus cinnabarinus GN=BN946_scf184888.g25 PE=4 SV=1 - [A0A060SNK6_PYCCI] | 1,93 | 0,74 | 1 | 1 | 1 | 1 | 1081 | 120,4 | 6,40 |
| A0A060STA7 | Uncharacterized protein OS=Pycnoporus cinnabarinus GN=BN946_scf184823.g8 PE=4 SV=1 - [A0A060STA7_PYCCI] | 1,91 | 1,57 | 1 | 1 | 1 | 1 | 508 | 57,6 | 5,39 |

| | | | | | | | | | | |
|---|---|---|---|---|---|---|---|---|---|---|
| A0A060SC37 | Glucose oxidase OS=Pycnoporus cinnabarinus GN=BN946_scf184803.g17 PE=3 SV=1 - [A0A060SC37_PYCCI] | 1,90 | 1,29 | 1 | 1 | 1 | 1 | 620 | 65,6 | 5,58 |
| A0A060SUM4 | Uncharacterized protein OS=Pycnoporus cinnabarinus GN=BN946_scf185034.g6 PE=4 SV=1 - [A0A060SUM4_PYCCI] | 0,00 | 7,51 | 1 | 1 | 1 | 1 | 466 | 51,8 | 6,11 |
