## Supplementary material for "Secretome characterization of the lignocellulose-degrading fungi *Pycnoporus sanguineus* and *Ganoderma applanatum*": Table S3

Table S3. Detailed information of 73 proteins identified from G. *applanatum* secretome

The original description of proteins highlighted in yellow was modified in Table 3 in order to mention the biological family which they belong to.

| Accession | Description | Score | Coverage | # Proteins | # Unique Peptides | # Peptides | # PSMs | # AAs | MW [kDa] | calc. pI |
|---|---|---|---|---|---|---|---|---|---|---|
| GL27550-R1_1 | glycoside hydrolase family 3 protein [Laccaria bicolor S238N-H82] gb\|EDR09330.1\| | 34,53 | 9,46 | 2 | 6 | 6 | 13 | 793 | 85,4 | 5,06 |
| GL23290-R1_1 | Probable beta-galactosidase C OS=Aspergillus niger (strain CBS 513.88 / FGSC A1513) GN=lacC PE=3 SV=1 | 33,87 | 5,05 | 1 | 5 | 6 | 13 | 1050 | 112,9 | 5,36 |
| GL27858-R1_1 | copper radical oxidase [Phanerochaete chrysosporium] | 31,28 | 6,51 | 1 | 4 | 4 | 12 | 922 | 98,4 | 4,49 |
| GL20779-R1_1 | peptidase M28 [Arthroderma otae CBS 113480] gb\|EEQ31398.1\| | 16,68 | 15,06 | 1 | 5 | 5 | 6 | 498 | 52,8 | 5,06 |
| GL30909-R1_1 | alpha-galactosidase [Phanerochaete chrysosporium] | 14,91 | 11,93 | 1 | 4 | 4 | 5 | 436 | 47,4 | 5,55 |
| GL22886-R1_1 | beta-xylosidase [Postia placenta Mad-698-R] gb\|EED80923.1\| | 11,91 | 5,48 | 2 | 3 | 4 | 5 | 803 | 86,7 | 5,21 |
| GL29240-R1_1 | Serine-type carboxypeptidase F OS=Aspergillus niger GN=pepF PE=1 SV=3 | 10,39 | 4,25 | 1 | 2 | 2 | 4 | 518 | 57,1 | 4,83 |
| GL23422-R1_1 | glycoside hydrolase family 92 protein [Schizophyllum commune H4-8] gb\|EFJ03492.1\| | 10,39 | 3,86 | 1 | 3 | 3 | 4 | 854 | 93,1 | 5,24 |
| GL16398-R1_1 | Laccase OS=Pycnoporus cinnabarinus GN=LCC3-1 PE=1 SV=1 | 10,38 | 5,77 | 1 | 2 | 2 | 3 | 520 | 56,6 | 4,87 |
| GL23225-R1_1 | Probable beta-galactosidase A OS=Neosartorya fischeri (strain ATCC 1020 / DSM 3700 / FGSC A1164 / NRRL 181) GN=lacA PE=3 SV=1 | 9,88 | 1,54 | 2 | 2 | 3 | 4 | 2072 | 223,8 | 5,12 |
| GL29486-R1_1 | Laccase OS=Pycnoporus cinnabarinus GN=LCC3-1 PE=1 SV=1 | 8,29 | 2,50 | 1 | 2 | 2 | 3 | 520 | 56,2 | 6,04 |
| GL20521-R1_1 | Putative amidase C869.01 OS=Schizosaccharomyces pombe (strain ATCC 38366 / 972) GN=SPAC869.01 PE=2 SV=1 | 8,11 | 5,26 | 1 | 2 | 2 | 3 | 551 | 57,5 | 5,01 |
| GL15069-R1_1 | hypothetical protein SCHCODRAFT_103662 [Schizophyllum commune H4-8] gb\|EFJ03568.1\| | 8,04 | 4,61 | 1 | 1 | 1 | 3 | 347 | 36,7 | 4,75 |
| GL24962-R2_1 | Histone H4 OS=Phanerochaete chrysosporium GN=H4.1 PE=3 SV=2 | 6,76 | 8,20 | 5 | 1 | 1 | 2 | 122 | 13,6 | 11,60 |
| GL27365-R1_1 | hypothetical protein SCHCODRAFT_231738 [Schizophyllum commune H4-8] gb\|EFJ03963.1\| | 5,70 | 6,79 | 1 | 1 | 1 | 2 | 221 | 23,0 | 7,03 |
| GL20532-R1_1 | mannose-6-phosphatase [Phanerochaete chrysosporium] | 5,42 | 7,25 | 1 | 1 | 2 | 2 | 345 | 37,7 | 5,11 |
| GL24763-R1_1 | mannose-6-phosphatase [Phanerochaete chrysosporium] | 5,34 | 6,30 | 2 | 1 | 2 | 2 | 365 | 39,7 | 6,80 |
| GL22158-R1_1 | GL22158-R1_1 | 5,31 | 7,41 | 1 | 2 | 2 | 2 | 189 | 20,1 | 10,17 |
| GL23249-R1_1 | Probable beta-galactosidase C OS=Aspergillus niger (strain CBS 513.88 / FGSC A1513) GN=lacC PE=3 SV=1 | 5,07 | 1,01 | 2 | 1 | 1 | 2 | 987 | 107,0 | 6,37 |

| | | | | | | | | | | |
|---|---|---|---|---|---|---|---|---|---|---|
| GL20904-R1_1 | hypothetical protein SCHCODRAFT_258939 [Schizophyllum commune H4-8] gb|EFI91349.1| | 4,63 | 2,16 | 2 | 2 | 2 | 2 | 1341 | 148,8 | 5,31 |
| GL26629-R1_1 | Actin-1 [Schizophyllum commune H4-8] sp|Q9Y702|ACT1_SCHCO RecName: Full=Actin-1; AltName: Full=Beta-actin gb|AAD38853.1|AF156157_1 actin 1 [Schizophyllum commune] dbj|BAA77815.1| beta-actin [Schizophyllum commune] gb|EFI91247.1| Actin-1 [Schizophyllum commune H4-8] | 4,46 | 9,12 | 1 | 2 | 2 | 2 | 373 | 41,5 | 5,68 |
| GL29943-R1_1 | translation elongation factor 1a [Schizophyllum commune H4-8] gb|EFJ02359.1| | 4,45 | 1,73 | 1 | 2 | 2 | 2 | 1101 | 119,4 | 8,53 |
| GL16765-R1_1 | Lipase 1 OS=Candida rugosa GN=LIP1 PE=1 SV=3 | 4,32 | 2,01 | 2 | 1 | 1 | 2 | 547 | 58,4 | 4,75 |
| GL19093-R1_1 | beta-xylosidase [Postia placenta Mad-698-R] gb|EED80923.1| | 4,31 | 2,23 | 2 | 1 | 2 | 2 | 808 | 87,5 | 5,21 |
| GL30117-R1_1 | hypothetical protein POSPLDRAFT_134924 [Postia placenta Mad-698-R] gb|EED79966.1| | 4,18 | 2,28 | 1 | 2 | 2 | 2 | 963 | 105,1 | 4,83 |
| GL31420-R1_1 | endopeptidase [Coprinopsis cinerea okayama7#130] gb|EAU84813.1| | 3,99 | 3,72 | 1 | 2 | 2 | 2 | 430 | 46,5 | 5,01 |
| GL18722-R1_1 | ATP synthase subunit alpha, mitochondrial OS=Neurospora crassa (strain ATCC 24698 / 74-OR23-1A / CBS 708.71 / DSM 1257 / FGSC 987) GN=atp-1 PE=3 SV=1 | 3,97 | 1,24 | 1 | 1 | 1 | 2 | 566 | 61,7 | 9,06 |
| GL29490-R1_1 | Laccase-2 OS=Trametes villosa GN=LCC2 PE=3 SV=1 | 3,68 | 2,95 | 1 | 1 | 1 | 1 | 509 | 54,4 | 4,93 |
| GL24346-R1_1 | N-acetylhexosaminidase [Postia placenta Mad-698-R] gb|EED78592.1| | 3,31 | 2,87 | 2 | 1 | 1 | 1 | 557 | 60,0 | 5,08 |
| GL30358-R1_1 | tRNA (adenine-N(1)-)-methyltransferase non-catalytic subunit TRM6 OS=Cryptococcus neoformans var. neoformans serotype D (strain JEC21) GN=TRM6 PE=3 SV=1 | 3,21 | 8,13 | 1 | 1 | 1 | 2 | 492 | 54,5 | 6,15 |
| GL16281-R1_1 | d-4,5 unsaturated -glucuronyl hydrolase-like protein [Postia placenta Mad-698-R] gb|EED77542.1| | 3,07 | 1,97 | 1 | 1 | 1 | 1 | 406 | 44,3 | 5,17 |
| GL30723-R1_1 | Uncharacterized oxidoreductase C23D3.11 OS=Schizosaccharomyces pombe (strain ATCC 38366 / 972) GN=SPAC23D3.11 PE=2 SV=2 | 3,02 | 10,84 | 1 | 1 | 1 | 1 | 286 | 30,9 | 8,22 |
| GL25498-R1_1 | hypothetical protein SCHCODRAFT_69684 [Schizophyllum commune H4-8] gb|EFI94947.1| | 2,99 | 5,38 | 1 | 1 | 1 | 1 | 372 | 37,9 | 4,83 |
| GL21099-R1_1 | glycoside hydrolase family 95 protein [Laccaria bicolor S238N-H82] gb|EDQ99849.1| | 2,83 | 1,32 | 1 | 1 | 1 | 1 | 831 | 90,2 | 5,10 |
| GL23044-R2_1 | GL23044-R2_1 | 2,77 | 5,85 | 2 | 1 | 1 | 1 | 188 | 20,6 | 6,20 |

| | | | | | | | | | | |
|---|---|---|---|---|---|---|---|---|---|---|
| GL24688-R1_1 | RecName: Full=Heat shock protein HSS1 gb\|AAB93665.1\| HSS1 [Puccinia graminis f. sp. tritici] | 2,72 | 1,80 | 1 | 1 | 1 | 1 | 612 | 66,7 | 5,30 |
| GL30143-R1_1 | proteasome subunit alpha type 5 [Coprinopsis cinerea okayama7#130] gb\|EAU92402.1\| | 2,71 | 3,83 | 1 | 1 | 1 | 1 | 261 | 29,2 | 5,73 |
| GL19832-R1_1 | GL19832-R1_1 | 2,67 | 17,39 | 1 | 1 | 1 | 1 | 138 | 15,8 | 11,49 |
| GL20524-R1_1 | Polyporopepsin OS=Irpex lacteus PE=1 SV=1 | 2,65 | 2,17 | 2 | 1 | 1 | 1 | 415 | 43,8 | 5,40 |
| GL27154-R1_1 | GL27154-R1_1 | 2,62 | 4,76 | 1 | 1 | 1 | 1 | 273 | 27,6 | 6,70 |
| GL20698-R2_1 | Probable mannosyl-oligosaccharide alpha-1,2-mannosidase 1B OS=Aspergillus terreus (strain NIH 2624 / FGSC A1156) GN=mns1B PE=3 SV=1 | 2,61 | 2,17 | 2 | 1 | 1 | 1 | 507 | 55,6 | 5,08 |
| GL22350-R1_1 | GL22350-R1_1 | 2,60 | 5,20 | 1 | 1 | 1 | 1 | 442 | 50,0 | 5,34 |
| GL29484-R1_1 | Ribosomal RNA processing protein 1 homolog B OS=Homo sapiens GN=RRP1B PE=1 SV=3 | 2,58 | 5,88 | 1 | 1 | 1 | 1 | 357 | 39,6 | 7,97 |
| GL18249-R1_1 | glycoside hydrolase family 92 protein [Schizophyllum commune H4-8] gb\|EFI98309.1\| | 2,51 | 1,86 | 1 | 1 | 1 | 1 | 862 | 92,7 | 4,97 |
| GL21683-R1_1 | Probable glucan endo-1,6-beta-glucosidase B OS=Aspergillus terreus (strain NIH 2624 / FGSC A1156) GN=exgB PE=3 SV=1 | 2,49 | 2,46 | 1 | 1 | 1 | 1 | 487 | 53,6 | 5,49 |
| GL28315-R1_1 | Vegetative incompatibility protein HET-E-1 OS=Podospora anserina (strain S / DSM 980 / FGSC 10383) GN=HET-E1 PE=4 SV=1 | 2,38 | 5,47 | 1 | 1 | 1 | 1 | 567 | 63,2 | 5,54 |
| GL30351-R1_1 | cellobiohydrolaseI [Polyporus arcularius] | 2,37 | 1,97 | 3 | 1 | 1 | 1 | 457 | 49,1 | 4,65 |
| GL18873-R1_1 | GL18873-R1_1 | 2,31 | 3,14 | 1 | 1 | 1 | 1 | 350 | 37,0 | 4,79 |
| GL27736-R1_1 | GL27736-R1_1 | 2,27 | 3,27 | 1 | 1 | 1 | 1 | 336 | 37,7 | 8,10 |
| GL21313-R1_1 | glyceraldehyde-3-phosphate dehydrogenase [Ganoderma lucidum] gb\|ABD64598.1\| | 2,25 | 2,82 | 2 | 1 | 1 | 1 | 319 | 34,3 | 6,39 |
| GL24652-R1_1 | predicted protein [Laccaria bicolor S238N-H82] gb\|EDR14338.1\| | 2,24 | 2,22 | 1 | 1 | 1 | 1 | 541 | 57,9 | 5,78 |
| GL22859-R1_1 | Probable alpha/beta-glucosidase agdC OS=Neosartorya fumigata (strain ATCC MYA-4609 / Af293 / CBS 101355 / FGSC A1100) GN=agdC PE=3 SV=1 | 2,21 | 1,12 | 1 | 1 | 1 | 1 | 891 | 97,8 | 6,62 |
| GL23580-R1_1 | glycoside hydrolase family 15 protein [Laccaria bicolor S238N-H82] gb\|EDR13753.1\| | 2,15 | 1,39 | 1 | 1 | 1 | 1 | 574 | 61,0 | 5,27 |
| GL29873-R2_1 | GL29873-R2_1 | 2,14 | 1,07 | 2 | 1 | 1 | 1 | 838 | 87,9 | 5,07 |
| GL21592-R1_1 | Aspartyl-tRNA synthetase, cytoplasmic OS=Schizosaccharomyces pombe (strain ATCC 38366 / 972) GN=dps1 PE=1 SV=1 | 2,11 | 1,92 | 2 | 1 | 1 | 1 | 520 | 58,5 | 6,77 |
| GL30540-R1_1 | glycoside hydrolase family 74 [Phanerochaete chrysosporium] | 2,09 | 1,11 | 1 | 1 | 1 | 1 | 722 | 76,2 | 4,93 |

| | | | | | | | | | | |
|---|---|---|---|---|---|---|---|---|---|---|
| GL22147-R1_1 | hypothetical protein POSPLDRAFT_107968 [Postia placenta Mad-698-R] gb\|EED80026.1\| | 2,08 | 2,64 | 1 | 1 | 1 | 1 | 455 | 49,4 | 5,36 |
| GL25209-R1_1 | GL25209-R1_1 | 2,03 | 2,94 | 1 | 1 | 1 | 1 | 306 | 33,4 | 7,55 |
| GL22710-R1_1 | GL22710-R1_1 | 2,02 | 2,05 | 1 | 1 | 1 | 1 | 438 | 46,3 | 4,53 |
| GL17521-R1_1 | GL17521-R1_1 | 2,00 | 1,99 | 1 | 1 | 1 | 1 | 452 | 51,5 | 7,36 |
| GL18489-R1_1 | hypothetical protein SCHCODRAFT_45589 [Schizophyllum commune H4-8] gb\|EFJ02307.1\| | 2,00 | 0,97 | 1 | 1 | 1 | 1 | 924 | 100,9 | 5,06 |
| GL20757-R1_1 | predicted protein [Laccaria bicolor S238N-H82] gb\|EDR10174.1\| | 2,00 | 3,97 | 1 | 1 | 1 | 1 | 353 | 37,6 | 4,55 |
| GL20594-R1_1 | hypothetical protein SCHCODRAFT_107709 [Schizophyllum commune H4-8] gb\|EFI97631.1\| | 1,98 | 1,80 | 1 | 1 | 1 | 1 | 555 | 58,8 | 5,85 |
| GL21504-R1_1 | Uncharacterized protein PB2B2.06c OS=Schizosaccharomyces pombe (strain ATCC 38366 / 972) GN=SPBPB2B2.06c PE=2 SV=1 | 1,97 | 1,25 | 1 | 1 | 1 | 1 | 640 | 70,9 | 5,47 |
| GL19409-R1_1 | hypothetical protein CC1G_03004 [Coprinopsis cinerea okayama7#130] gb\|EAU85981.2\| | 1,97 | 0,36 | 1 | 1 | 1 | 1 | 1968 | 219,2 | 5,40 |
| GL25858-R1_1 | N-acetyltransferase eso1 OS=Schizosaccharomyces pombe (strain ATCC 38366 / 972) GN=eso1 PE=1 SV=1 | 1,95 | 0,81 | 1 | 1 | 1 | 1 | 745 | 82,4 | 6,70 |
| GL23600-R1_1 | glycoside hydrolase family 15 protein [Laccaria bicolor S238N-H82] gb\|EDR13753.1\| | 1,95 | 1,29 | 1 | 1 | 1 | 1 | 541 | 57,3 | 4,45 |
| GL22664-R1_1 | GL22664-R1_1 | 1,95 | 1,99 | 1 | 1 | 1 | 1 | 702 | 76,2 | 7,17 |
| GL22513-R1_1 | Histone H2A OS=Agaricus bisporus PE=2 SV=3 | 1,93 | 6,52 | 4 | 1 | 1 | 1 | 138 | 14,6 | 10,08 |
| GL16919-R1_1 | GL16919-R1_1 | 1,92 | 4,18 | 1 | 1 | 1 | 1 | 383 | 41,0 | 5,54 |
| GL30320-R1_1 | Elongation factor Tu, mitochondrial OS=Schizosaccharomyces pombe (strain ATCC 38366 / 972) GN=tuf1 PE=2 SV=1 | 1,90 | 1,45 | 1 | 1 | 1 | 1 | 482 | 52,4 | 8,31 |
| GL23555-R2_1 | Potassium voltage-gated channel subfamily D member 3 OS=Oryctolagus cuniculus GN=KCND3 PE=2 SV=1 | 1,90 | 1,79 | 2 | 1 | 1 | 1 | 390 | 43,3 | 5,30 |
| GL30052-R1_1 | GL30052-R1_1 | 0,00 | 9,29 | 1 | 1 | 1 | 1 | 280 | 30,8 | 5,54 |
